# Targeting of a STING Agonist to Perivascular Macrophages in Prostate Tumors Delays Resistance to Androgen Deprivation Therapy

**DOI:** 10.1101/2024.04.11.589003

**Authors:** Haider Al-janabi, Katy Moyes, Richard Allen, Matthew Fisher, Mateus Crespo, Bora Gurel, Pasquale Rescigno, Johann De Bono, Harry Nunns, Christopher Bailey, Anna Juncker-Jensen, Munitta Muthana, Wayne A Phillips, Helen B Pearson, Mary Ellen-Taplin, Janet E. Brown, Claire E Lewis

**Affiliations:** Division of Clinical Medicine, University of Sheffield Medical School, Beech Hill Rd, Sheffield, S10 2RX, UK; The Institute of Cancer Research & The Royal Marsden Hospital, Sutton, Surrey, UK; Neogenomics Laboratories, 31 Columbia, Aliso Viejo, CA 92656, USA; LipExoGen Biotech, 101 N. Haven Street, Baltimore, MD21224. USA; The Peter MacCallum Cancer Centre, Melbourne, Victoria, Australia; The European Cancer Stem Cell Research Institute, School of Biosciences, Cardiff University Maindy Road, Cardiff, CF24 4HQ. UK; Dana-Farber Cancer Institute, 450 Brookline Ave Boston MA 02215, USA

**Author notes:** Corresponding Author: Claire E. Lewis, Division of Clinical Medicine, University of Sheffield Medical School, Beech Hill Rd., Sheffield, S10 2RX, UK.

**Keywords:** Prostate cancer, ADT, Immunotherapy, Lipid nanoparticles, cGAMP, STING, Perivascular macrophages, CRPC

## Abstract

**Background:** Androgen deprivation therapy (ADT) is a frontline treatment for prostate cancer but often leads to the development of castration resistant prostate cancer (CRPC). This causes tumors to regrow and metastasize, despite ongoing treatment, and impacts negatively on patient survival. ADT is known to stimulate the accumulation of immunosuppressive cells like protumoral tumor-associated macrophages (TAMs), myeloid-derived suppressor cells and regulatory T cells in prostate tumors, as well as hypofunctional T cells. Protumoral TAMs have been shown to accumulate around tumor blood vessels during chemotherapy and radiotherapy, where they drive tumor relapse. Our aim was to see if such perivascular (PV) TAMs accumulated in ADT-treated prostate tumors prior to CRPC, and, if so, to selectively target these PV cells with a potent immunostimulant, interferon beta (IFNβ), an attempt to stimulate anti-tumor immunity and delay CRPC.

**Methods:** We first used quantitative, multiplex immunofluorescence to assess the effects of ADT on distribution and activation status of TAMs, CD4+ T cells, CD8+ T cells and NK cells in mouse and human prostate tumors. We then used antibody-coated, lipid nanoparticles to selectively target a STING agonist, 2′3′-cGAMP (cGAMP), to PV TAMs in mouse prostate tumors during ADT.

**Results:** TAMs accumulated at high density around blood vessels in ADT-treated primary mouse and human prostate tumors prior to CRPC, where they expressed markers of a protumoral phenotype, folate receptor beta (FRβ), MRC1 (CD206), SIGLEC1 (CD169) and VISTA. Additionally, higher numbers of inactive (PD-1-) CD8+ T cells and reduced numbers of active (CD69+) NK cells were also present in PV tumor areas after ADT. LNPs coated with antibody to FRβ selectively delivered cGAMP to PV TAMs in ADT-treated tumors where they activated STING and expression of IFNβ by these cells. This resulted in a marked increase in the density of active CD4+ T cells, CD8+T cells and NK cells in PV tumor areas, and significantly delayed in the onset of CRPC.

**Conclusion:** Together, our data indicate that targeting a STING agonist to PV TAMs could be used to extend the treatment window for ADT in prostate cancer.

**KEY MESSAGES:** *What is already known about the topic:* Androgen deprivation therapy (ADT) is a frontline treatment for prostate cancer. However, tumors often develop resistance and start to regrow and metastasize – a condition called castration resistance prostate cancer (CRPC). Prostate cancer is considered to be an immunologically ‘cold’ tumor type and while ADT stimulates tumor infiltration by cytotoxic (CD8+) T cells, they are largely hypofunctional, possibly due to the immunosuppressive tumor microenvironment.

*What this study adds:* This study is the first to demonstrate that FRβ+ macrophages with a immunosuppressive phenotype accumulate around blood vessels in mouse and human prostate tumors during ADT, prior to the onset of CRPC. Lipid nanoparticles coated with an antibody to FRβ+ were then used to deliver a STING agonist selectively to these perivascular (PV) cells during ADT. This triggered STING signalling and the release of the potent immunostimulant, interferon beta, by PV macrophages, which then activated tumour-infiltrating CD4+ and CD8+ T cells, and delayed the onset of CRPC.

*How this study might affect research, practice or policy:* The delivery of an immunostimulant specifically to PV regions of tumors represents a new, more targeted form of immunotherapy that ensures the activation of T cells as soon as they cross the vasculature into tumors. This new approach could be used to extend the treatment window for neoadjuvant ADT in men with localised prostate tumors. In doing so, it would delay/circumvent the need for additional treatments like radiotherapy and/or or prostatectomy.

## INTRODUCTION

Prostate cancer is the second most common cancer in men. Androgens are known to stimulate the progression of this disease via androgen receptors (ARs) over-expressed on cancer cells. This prompted the development of drugs that their either reduce/ablate androgen synthesis in the testes or reduce AR signalling. Collectively, these anti-androgens are known as androgen deprivation therapy (ADT).^1^

Men with intermediate/high-risk, localised prostate cancer are usually offered neoadjuvant ADT. Although this initially reduces tumor burden, it is often not curative and tumors develop resistance to ADT, a condition called castration resistant prostate cancer (CRPC), which then metastasises to local lymph nodes and other tissues. Despite the development of various treatment options for CRPC, including androgen/AR signaling inhibitors such as abiraterone and enzalutamide, and PARP inhibitors, the median survival rate for men with metastatic CRPC remains poor at under three years.^2–4^ This highlights the unmet clinical need for novel therapies that can prevent/delay the emergence of CRPC.

Mechanistically, the emergence of CRPC has been strongly linked to aberrant AR signalling, defective DNA damage/repair pathways, and the intratumoral synthesis of androgens^3^. However, AR-independent mechanisms can also facilitate CRPC, including loss of function mutations in the tumor suppressor genes, p53 and PTEN, and deregulation of the FGF, TGFβ, RAS/MAPK, hedgehog and Wnt/β- catenin signaling pathways. Although therapeutic agents that target one or more of the above have shown promise in the clinic, outcomes remain poor^4^.

ADT has been shown to increase the frequency of various immune effector cells in both mouse and human prostate tumors.^5–8^ For example, several reports have shown that tumor infiltration by CD8+ Τ cells increases with ADT, although this fails to impact favourably on relapse or survival as these cells have reduced cytotoxic function.^5–8^ Additionally, ADT has been shown to impair T cell priming by antigen-presenting cells and T cell expression of cytotoxicity-related genes (granzymes and perforins).^5,9–11^ It is also reported to increase the intratumoral abundance of various immunosuppressor cell types including tumor-associated macrophages (TAMs), myeloid-derived suppressor cells (MDSCs), and regulatory T cells (Tregs).^5,12–14^

Agonists of stimulator of interferon genes (STING)-signalling pathway in cells are emerging as an exciting form of immunotherapy for cancer.^15,16^ These include factors that activate STING like cyclic guanosine monophosphate (GMP)–adenosine monophosphate (AMP) synthase (cGAS) or cyclic dinucleotide 2′3′-cGAMP (cGAMP).^17^ Activation of STING in cells causes them to upregulate type I interferons (IFNs), including IFNα and INFβ.^18^ In tumors, the latter have been shown to stimulate anti-tumor immunity via multiple mechanisms including the cross-priming and activation of CD8+ T cells by antigen presenting cells, as well as the activation of both CD4+ T cells and NK cells.^19,20^ However, the clinical efficacy STING agonists of administered systemically is compromised by their rapid excretion, low bioavailability, lack of specificity, and adverse, off-target, side effects^21^. Furthermore, intratumoral administration is limited by the accessibility of tumors. To overcome these limitations, a number of delivery systems have been used to provide protection for STING agonists in the circulation and enable them to access tumors. These include their encapsulation in lipid nanoparticles (LNPs), exosomes and bacterial vectors^22^.

In the present study, we show that ADT causes a marked change in the immune landscape immediately adjacent to blood vessels in both mouse and human prostate tumors just prior to the onset of CRPC. For example, a marked increase in the density of pro-tumoral TAMs and naïve (PD-1-) CD8+ T cells was seen in these perivascular (PV) areas, indicating the ability of ADT to promote an immunosuppressive, PV niche.

We reasoned that the systemic targeting of a STING agonist LNPs to such protumoral, PV TAMs would be effective due to their close proximity to tumor blood vessels. So, we synthesised LNPs containing cGAMP and coated them with an antibody raised against a receptor we show is expressed by protumoral PV TAMs in ADT-treated tumors (folate receptor beta). Following systemic administration of these LNPs to ADT-treated mice bearing prostate tumors, cGAMP was selectively delivered to PV TAMs, triggering STING signaling and their upregulation of IFNβ. This resulted in the activation of tumor-infiltrating immune effectors like T and NK cells and delayed the onset of CRPC.

## METHODS

### Orthotopic mouse models of primary prostate cancer

#### Myc-CaP implants

Myc-CaP cells^23^ stably expressing luciferase were cultured in Dulbecco’s Modified Eagle Medium (DMEM) supplemented with 10% fetal bovine serum (FBS) and 1% penicillin/streptomycin (P/S) and were tested regularly for mycoplasma. 50,000 Myc-CaP cells (1:1 PBS:Matrigel) were injected into the dorsal prostate lobe of male FVB mice (aged 6-8 weeks). Mice with detectable tumors 7 days later were randomised into experimental groups and injected subcutaneously (once at day 7 alone or with a second dose on day 14) with either vehicle control (PBS) or degarelix (25 mg/kg) (Ferring Pharmaceuticals Inc., Parsippany, NJ). (n=5-8/treatment arm). Tumor growth was then monitored every 2-3 days using an IVIS Spectrum imaging system (ie. 30 mins after injection with 90 mg/kg d-luciferin).

All Myc-CaP experiments were carried out in compliance with UK Home Office Regulations as specified in the Animals (Scientific Procedures) Act of 1986, and were approved by the Animal Welfare and Ethical Review Body of the University of Sheffield.

#### Transgenic Pten-deficient mice

*PBiCre^+/-^;Pten^fl/fl^* mice were generated as described previously.^24^ At 200 days of age, mice were randomly assigned to sham-castrated or castrated groups. Prostate tumors were harvested 2 weeks after surgery, snap frozen in liquid nitrogen, and OCT-embedded before being cryosectioned.

Transgenic mouse experiments adhered to the guidelines outlined in the National Health and Medical Research Council (NHMRC) Australian Code of Practice for the Care and Use of Animals for Scientific Purposes andwere approved by the Animal Experimentation Ethics Committee at the Peter MacCallum Cancer Centre.

### Quantitative immunofluorescence staining of mouse prostate tumors

Cryosections of mouse prostate tumors were fixed with ice cold acetone for 10 min, blocked with 5% goat serum and 10% mouse FcR blocking solution before being incubated for 40 mins with primary antibodies; F4/80 (BIO-RAD, 1:100), FRβ (BioLegend, 1:100), CD169 (BioLegend, 1:100), VISTA (BioLegend, 1:100), CD206 (MRC1) (BioLegend, 1:100), CD31 (BioLegend, 1:100), CD8 (BioLegend, 1:100), PD-1 (BioLegend, 1:100), phopho-STING (ThermoFisher 1:100), NK-1.1 (Biolegend 1:100), CD69 (Biolegend 1:100), CD4 (Biolegend 1:100), and IFNbeta (Invitrogen, 1:100). Primary rabbit antibodies were detected using goat anti-rabbit (IgG) Alexa Fluor 555 (Fisher Scientific 1:400). All sections were counter-stained with 50ng/ml DAPI solution before washing and mounting.

A Nikon A1 confocal microscope was used to capture 5 randomly selected images/tumor (x20 magnification). The acquired images were subsequently analyzed using QuPath (version 0.4.3) or ImageJ. For analysis of cell density relative to blood vessels, cells <15 µm from CD31+ blood vessels were defined as PV. These were counted in each ROI and divided by its vessel area to estimate the PV density/ROI. Cells >15 µm from CD31+ blood vessels were defined as non-PV, the density of which was calculated by dividing cell numbers in these areas of a given ROI, by the total non-PV area of that ROI.

### Human primary prostate tumors

Sections were cut from localized, human prostate tumors removed from men in the following two groups:

1. Matched pre-treatment biopsies and post-treatment, radical prostatectomies (RPs) from twenty patients who received neoadjuvant ADT for six months at the Dana-Farber Cancer Institute, Boston, USA. Additionally, pathological staging of prostatectomy samples (ie. after ADT) enabled us to divide these into ‘responder’ or ‘non-responder’ groups. Responders were defined as patients with residual prostate cancer of </= 5mm, T stage = 0-2, after ADT. Whereas non-responders had tumors that were T stage >/= 3a after ADT (and so were starting to regrow and show signs of CRPC). There was no regional lymph node involvement in either group.
2. Six ‘untreated’ tumors (ie. not given ADT; 4 prostatectomies and 2 transurethral resections of the prostate *or ‘*TURPs’), and five TURPs from patients who received ADT (median treatment time, 7 months). Three patients in the latter group had draining lymph node involvement (ie. early metastasis). Both groups of samples were supplied by The Institute of Cancer Research, London.

Tumor group 1 (above) was used to assess the co-expression of CD68 and FRβ in PV versus non-PV tumor areas in <u>matched</u> tumor samples before and after ADT. An antibody to CD31 was also used to label blood vessels. Antigen retrieval was performed using a pressure cooker in conjunction with Dako 10X retrieval solution (S1699). Subsequently, the sections were blocked to reduce non-specific binding using 10% goat serum and 1% TBS (at room temperature for 30 minutes).

### Quantitative immunofluorescence staining of human prostate tumors

For tumor group 1, sections were incubated in primary antibodies diluted in 1% bovine serum albumen (BSA) overnight at 4°C, then washed x3 in TBS, followed by secondary antibodies at room temperature for 1h and DAPI for 3 minutes, before washing and mounting. The primary antibodies used were a mouse monoclonal (IgG3) anti-human CD68 (Dako, Clone PG-M1 1:200), a mouse monoclonal (IgG1) anti-CD31 (Antibodies.com, Clone JC/70A 1:100) and a rabbit polyclonal anti-FRβ (Abcam 1:300). The negative controls for these primary antibodies were species, isotype and concentration-matched antibodies. The secondary antibodies were a goat anti-mouse IgG3-Alexa Fluor 488 (Fisher Scientific 1:200), a goat anti-mouse IgG1-Alexa Fluor 647 (ThermoFisher 1:200) and a goat anti-rabbit IgG-Alexa Fluor 555 (Fisher Scientific 1:100).

For tumor group 2, the ‘MultiOmyx’ procedure was used to detect CD8, PD-1 and CD31 as described by us previously^26^ in untreated and ADT-treated tumors. The antibodies used were mouse anti-human CD8 (Dako), a rabbit anti-human PD-1 (Abcam), and a mouse anti-human CD31/PECAM-1 (Cell Signaling). All three were conjugated to fluorophores and diluted with 3% (wt/vol) BSA (to working concentrations optimized previously^26^). They were applied to sections for 1h at RT, then washed in PBS and high-resolution images collected from 20 regions of interest (ROIs) across viable tumor areas using a 20x objective on an INCell analyzer 2200 microscope (GE Healthcare Life Sciences). An AI-based, advanced analytics platform, proprietary to NeoGenomics Labs, called ‘NeoLYTX™’, was used to quantify and analyse subsets of PD-1- and PD-1+CD8+ T cells, and CD31+ blood vessels in PV (<15 µm from CD31+ blood vessels) and non-PV (>15 µm from CD31+ blood vessels) areas of tumors.

### Generation of lipid nanoparticles to selectively target PV TAMs *in vivo*

LNPs containing the STING (<u>ST</u>imulator of <u>IF</u>N <u>G</u>enes) agonist, cGAMP, or an inert version of this molecule and coated with one of two antibodies, a rat anti-mouse FRβ or a rat IgG2a (isotype matched), control antibody (both from BioLegend) were synthesized by LipExoGen Biotech (**Suppl. Figure 9A**).

For this, 20mg/mL D-Lin-MC3-DMA (MC3, MedChemExpress) in ethanol, was mixed 1:1 (v/v) with 1,2-distearoyl-sn-glycero-3-phosphocholine (DSPC, Avanti Polar Lipids), cholesterol (Chol, Avanti Polar Lipids), 1,2-distearoyl-sn-glycero-3-phosphoethanolamine-N-[methoxy(polyethyleneglycol)- 2000] (DSPE-PEG2000, Avanti Polar Lipids) and OG488 DHPE (AAT Bioquest) in ethanol. The molar ratios of the components for the base formulation were as follows: MC3/Chol/DSPC/DSPE-PEG2000/DHPE-OG488 (50/38.5/9.8/1.5/0.2, mol/mol). cGAMP (cGAMP, InvivoGen, tlrl-nacga23) or its inactive control (InvivoGen, tlrl-nagpap) were dissolved in UltraPure water (Invitrogen) and diluted in acetate buffer, pH 4. LNPs were synthesized using a 3:1 aqueous:organic ratio and subsequently washed in UltraPure water using Amicon centrifugal columns (100 kDa). The encapsulation efficiency (EE%) of cGAMP (or its inactive control) were determined by HPLC.

Prior to the functionalization of the LNPs with antibodies, 1 mol% of DSPE-PEG2000 was post-inserted into LNPs. Fc-specific labelling (ie. to ensure antibody attachment to LNPs in the correct orientation for binding to FRβ on cells) was achieved by performing a click reaction between the DSPE-PEG2000-DBCO (BroadPharm) and terminal GlcNAz residues on the antibody carbohydrate domain. To obtain the latter, the SiteClick Antibody Azido Modification Kit (Invitrogen) was used to replace terminal galactose residues on the N-linked sugars in the Fc region with the azide-containing sugar GalNAz, which is reactive towards DBCO through strain-promoted alkyne-azide cycloaddition. The lipidated antibodies in DPBS(1X) were post-inserted into the LNPs according to the methods described by Swart and colleagues.^27^ Finally, an additional 2 mol% DSPE-PEG2000 was post-inserted into the LNPs via thin film hydration to minimize non-specific cellular uptake. The inclusion in LNPs of a phospholipid labelled on the head group with the bright, green-fluorescent fluorinated fluorescein dye (also called Oregon Green® 488), ‘DHPE-OG488’, will enable LNP uptake/distribution in tumors to be tracked by *in vivo*.

Mice bearing Myc-CaP tumors were assigned to one of eight treatment groups (5-8 mice/group): (i) Control – ie. PBS, alone); (ii) vehicle plus FRβ-antibody coated LNPs containing control cGAMP; (iii) vehicle plus FRβ-antibody-coated plus LNPs containing active cGAMP; (iv) (‘ADT’) degarelix as a single, sub.cut. dose of 25 mg/kg alone in PBS (Ferring Pharmaceuticals Inc., Parsippany, NJ); (v) degarelix plus FRβ-antibody coated LNPs containing inactive cGAMP (called ‘LNP(C)’ in some figures); (vi) degarelix plus FRβ-antibody coated LNPs containing active cGAMP (called ‘LNP(E)’ in some figures); (vii) degarelix plus control rat IgG-antibody coated LNPs containing inactive 2’3-cGAMP; (viii) degarelix plus control rat IgG-antibody coated LNPs containing active 2’3-cGAMP; (ix) degarelix plus LNPs with no antibody coating but containing active 2’3-cGAMP.

Mice received either a subcutaneous (s.c) injection of PBS alone or degarelix alone seven days after inoculation, or this followed by s.c. injections every two days with one of the LNP groups listed above. The mice were monitored daily for well-being and weighed every 2 days. Tumors were snap frozen prior to being embedded in optimum cutting temperature (OCT) compound for frozen sectioning.

### Flow cytometry

Myc-CaP cells were detached from 6-well plates using trypsin/EDTA, washed and re-suspended in the following primary antibodies: a rat monoclonal (IgG2a) anti-mouse FRβ (BioLegend,1:100) or a sheep polyclonal anti-mouse FRα (R&D Systems, 1:100) for 45-60 min. Cells were then washed twice and re-suspended in FACS buffer. Flow cytometry was performed using the BD LSR II flow cytometer and data processed using FlowJo Software.

### Statistical analysis

All data shown are means +/- SEMs. Dots on jitter plots represent values for individual tumors. All data were analyzed using GraphPad Prism 8.02 software. Data analysis was conducted blind and statistical analysis performed using the Mann-Whitney U-test with *p* values of < 0.05 considered to be significant.

## RESULTS

### ADT induces PV accumulation of protumoral TAMs in mouse and human prostate tumors

The effects of ADT on the distribution and phenoptype of TAMs were investigated just prior to CRPC in the immunocompetent Myc-CaP model. It caused an initial reduction in tumor burden within the first 7 days of treatment, but then started to regrow (ie. acquire CRPC) 7-10 days post-ADT treatment (**Figure 1A**). Sections from tumors harvested on day 17 from both control and ADT-treated mice were co-immunofluorescently stained for the macrophage marker F4/80 and the endothelial cell marker CD31. While both PV and non-PV F4/80+ TAMs were increased by ADT relative to the control group (**Figure 1B&C**), a significantly greater increase was seen in PV TAMs. Elevated PV F4/80+ TAMs were also observed prior to the acquisition of CRPC in a *Pten*-deficient transgenic mouse model of prostate cancer (*PBiCre^+/-^;Pten^fl/fl^*)^23^ 2-weeks post-surgical castration, compared to sham-castration (**Suppl. Figure 3A**).

**Figure 1.**
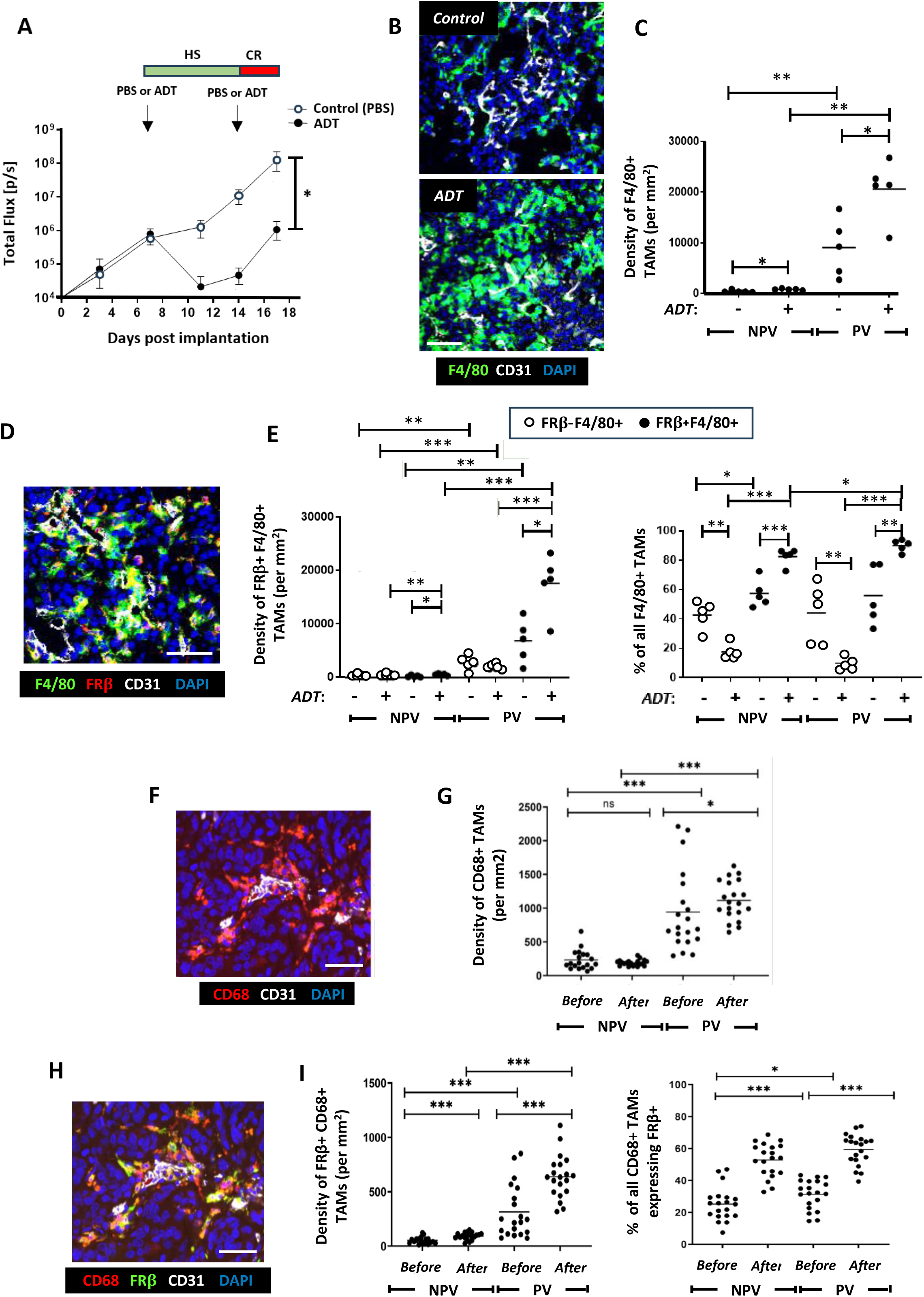
ADT stimulates the PV accumulation of FRβ+ TAMs in mouse (Myc-CaP) and localised, human prostate tumors. (**A**) Two phases of tumor response to the LHRH antagonist, degarelix, in Myc-CaP tumors: an initial, hormone sensitive (HS) period of tumor growth inhibition followed by the start of castration resistance (CR) when tumors start to regrow. In Myc-CaP tumors **(B-E)**, immunofluorescence staining shows that the density of both PV F4/80+ TAMs (**B&C**) - and the FRβ+ subset of these cells (**D-E**) increased by the start of CR in ADT-treated tumors. Similar changes occurred in non-PV tumor areas but to a lesser extent than in PV areas. The proportion of F4/80+ TAMs expressing FRβ also increased in PV and non-PV areas at this time. Immunofluorescence staining of matched human prostate tumors (**F&H**) sampled before and after ADT showed that ADT increased the PV density of PV CD68+ tumors (**G**) and the CD68+ TAM subset expressing FRβ (**I, left panel**). The proportion of CD68+ TAMs expressing FRβ rose in PV and non-PV tumor areas after ADT (**I, right panel**). (NPV = non-PV). Data are presented as means ± SEMs *p□<□0.05, **p□<□0.01, and ***p□<□0.001. Magnification bars = 50µm.

To determine if ADT also alters macrophage phenotype, we interrogated the phenotype of F4/80+ TAMs in PV and non-PV areas post-ADT using a panel of well-characterised markers of protumoral macrophages, including folate receptor beta (FRβ), CD169 (SIGLEC1), V-domain immunoglobulin suppressor of T *cell* activation (VISTA) and MRC1 (CD206), (**Figure 1D&E and Suppl. Figure 1**).^28–31^ Strikingly, the density of F4/80^+^ TAMs co-expressing each of the tumor-promoting cell surface macrophage markers analysed in PV regions were significantly upregulated upon ADT relative to the control. An increase in TAMs expressing these markers was also observed in non-PV regions but was significantly lower relative to than in PV regions (**Figure 1D&E** and **Suppl. Figure 1**). Collectively, these findings indicate that ADT promotes the accumulation of PV pro-tumor TAMs.

Given that both the densities and proportions of PV F4/80+TAMs expressing FRβ, CD169, VISTA or MRC1 were virtually identical (**Suppl. Figure 1 and 2A-C**), we investigated whether the same PV TAMs co-expressed these markers. Multiplex immunofluorescence staining for F4/80, FRβ, CD169 and VISTA confirmed that these markers were expressed by the same TAM subset. This showed that ADT treatment induces a subset of TAMs with a distinct protumoral phenotype in Myc-CaP tumors at the onset of CRPC, with significantly higher frequency in PV areas (**Suppl. Fig. 2D**).

**Figure 2.**
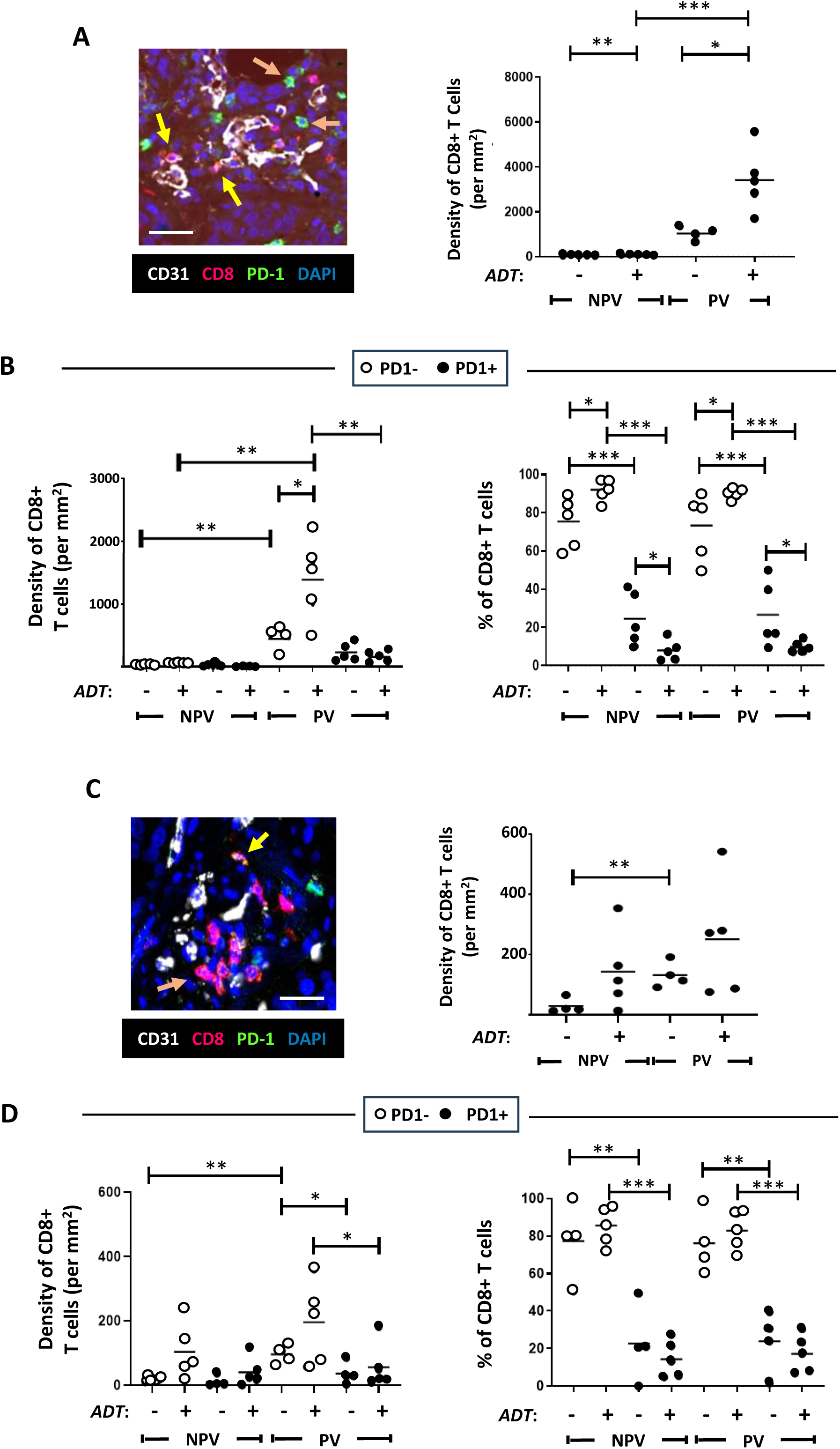
ADT stimulates the PV accumulation of PD-1-CD8+ T cells in mouse (Myc-CaP) (A,B) and human (C,D) prostate tumors. **(A,B).** (**A**) Representative fluorescence images showing the presence of mainly PD-1-CD8+ T cells in PV areas of ADT-treated Myc-CaP (**A**) and human (**C**) prostate tumors (left panels in both). In panels **A&C**, yellow arrows = PD-1+CD8+T cells, orange arrows = PD-1-CD8+T cells. ADT stimulates the PV accumulation of CD8+ T cells (**A&C**, right panels), which are mainly PD-1- (**B&D** left panels). The majority of CD8+ T cells lack expression of PD-1 across tumors, which increases further after ADT **(B,D**, right panels). (NPV = non-PV). Data are presented as means ± SEMs *p□<□0.05, **p□<□0.01, and ***p□<□0.001. Magnification bars = 50µm.

To establish if the observed induction of protumoral PV TAMs was specific to androgen deprivation in the Myc-CaP model, immunofluorescence staining was performed to co-localise F4/80 and MRC1 in primary prostate tumors from *PBiCre+;Pten^fl/fl^* mice, 2-weeks post-surgical castration. A similar increase in the PV density of MRC1+F4/80+ TAMs was observed (**Suppl. Figure 3A&B**). PV F4/80+ TAMs also expressed FRβ following castration (**Suppl. Figure 3**).

To investigate the clinical relevance of the above findings, we first examined matching human prostate tumor specimens collected before and after ADT for the human macrophage marker, CD68 and CD31. Immunofluorescence analysis revealed that while the density of CD68+ TAMs was significantly higher in PV than non-PV areas before and after ADT, their PV density increased further after ADT (**Figure 1F&G**).

The density of PV and non-PV FRβ+CD68+TAMs was significantly higher after ADT than before, but this effect of ADT was significantly greater in PV areas (**Figure 1H&I**).

We then investigated whether CD68+ TAM distribution correlated with tumor responses to ADT by dividing patients who received this treatment for six months into those that showed increased tumor growth during ADT (ie. were non-responders, ‘NRs’) or did not grow (ie. were responders, ‘Rs’). **Suppl**. **Figure 4A** there were no differences in CD68+ TAMs between Non-Rs and Rs, before and after ADT. While the FRβ+ subset of CD68+ TAMs showed a similar PV location to TAMs labelled for CD68 alone (ie both before and after ADT), the density of PV FRβ+CD68+ TAMs was significantly higher for NRs than R’s before ADT. There was a non-significant (p=0.07) trend for PV FRβ+CD68+ TAMs to also be higher in NRs than Rs after ADT (**Suppl**. **Figure 4B**). The non-PV density of these TAMs was also higher in NRs than Rs before ADT but this was significantly lower than in PV areas. (**Suppl**. **Figure 4B**). These data confirm the abundance of protumoral (FRβ+) TAMs in PV areas of human prostate tumors after ADT, especially those entering CRPC (ie. NRs).

### ADT alters the activation status of various immune effectors in PV areas

Given that protumoral TAMs were observed to increase in PV areas upon ADT, we reasoned that this might lead to (or coincide with) changes in effectors cells with cytotoxic potential (CD4+ and CD8+ T cells, and NK cells). To address this, we assessed the density and activation status of these immune effector cells in PV and non-PV areas of control and ADT-treated Myc-CaP tumors

We show that CD8+ T cell density dramatically increases significantly in PV areas after ADT (this was also observed in non-PV regions albeit in a significantly smaller CD8+ T cell population) (**Figure 2A**). Interestingly, our analysis of PD-1-expression by CD8+ T cells revealed that the majority (70%) of CD8+ T cells accumulating at PV regions upon ADT lacked this activation marker, and so were antigen-naïve (**Figure 2B**).

While CD8+ T cells tended towards a higher density in PV than non-PV areas of both untreated and ADT-treated tumors (**Figure 2C**, right panel), a significant increase in PD-1 negative CD8+ T cells was evident in PV (but not non-PV) areas after ADT (**Figure 2D**), resembling the Myc-CaP model. Together, these data indicate that ADT causes naive CD8+ T cells to accumulate in PV areas.

Analysis of CD4+ T cells in Myc-CaP prostate tumors showed that CD4+ T cells also principally accumulate in PV areas of both control and ADT-treated tumors (**Suppl. Fig. 5A**). However, in both PV and non-PV areas, the density and proportion of CD4+ T cells expressing PD-1 (approximately 50% in all groups) did not change during ADT (**Suppl. Fig. 5B)**. So, ADT did not alter the distribution or activation status of CD4+ T cells in Myc-CaP tumors.

NK cells were identified using the antibody, NK1.1, and their activation status investigated using CD69, an established marker of activation in NK cells (**Suppl. Fig. 5C&D**). NK cells were more frequent in PV than non-PV areas of both control and ADT-treated Myc-CaP tumors (**Suppl. Fig. 5C, right panel**), and were almost all CD69 positive (i.e. active). A similar trend was observed in non-PV areas, however the density of NK cells was significantly lower in these regions. Of note, differences in the PV density of CD69+NK cells in ADT-treated versus control tumors failed to reach significance (*p*=0.056) (**Suppl. Fig. 5D**).

### FR**β**-targeted lipid nanoparticles (LNPs) selectively target the STING agonist, cGAMP, to PV TAMs in ADT-treated Myc-CaP tumors, and delay CRPC

To take advantage of these immune-activating functions of the cGAMP-STING signaling pathway^18,19^ we generated LNPs containing cGAMP and targeted these to PV TAMs in ADT-treated tumors by coating them with an antibody to FRβ. The two main aims of this part of the study were to investigate whether FRβ-targeted LNPs could selectively deliver cGAMP to PV TAMs in ADT-treated tumors and stimulate them to express IFNβ, and whether this would stimulate anti-tumor immunity and delay the onset of CRPC.

As a prelude to the LNP *in vivo* experiment, we first confirmed that the FRβ antibody used to coat LNPs did not bind to a related molecule, FRα (known to be expressed by Myc-CaP cells). Flow cytometry and immunofluorescence staining using a specific FRα antibody alongside our FRβ antibody confirmed that Myc-CaP cells express FRα but not FRβ, and that the opposite was the case for TAMs in Myc-CaP tumors (**Suppl. Fig. 6B&C**).

Multiple, novel formulations of LNPs were synthesised containing either an active or inactive form of cGAMP, and coated with either an anti-mouse FRβ antibody or an IgG control antibody (**Figure 3A**; see method for LPN synthesis in **Suppl. Figure 6A**).

**Figure 3.**
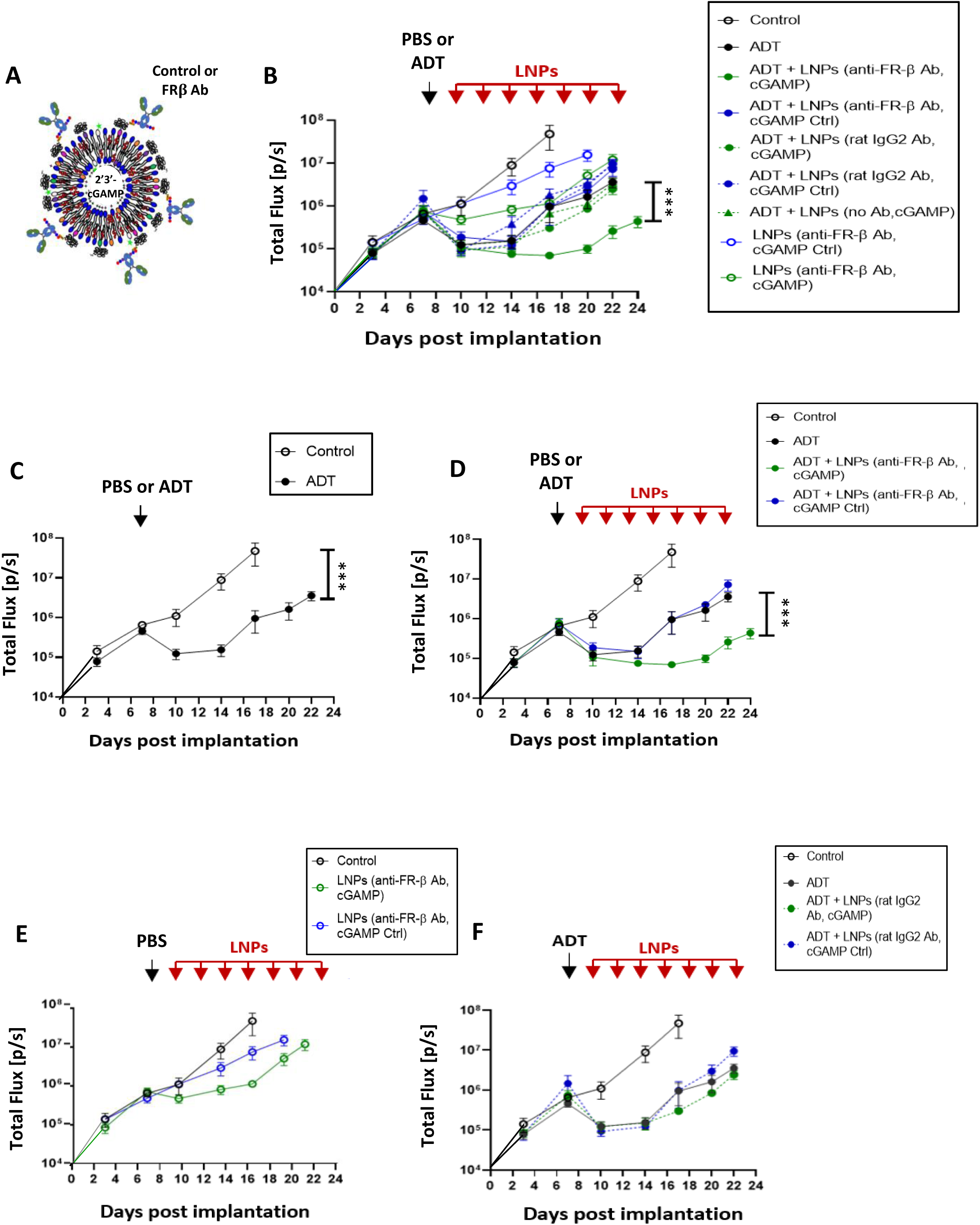
LNPs coated with FRβ-antibody target the STING agonist, cGAMP, to PV TAMs and delay CR in ADT-treated Myc-CaP tumors. (**A**) Design of LNPs used *in vivo*. The Fc regions of either a FRβ antibody or a control IgG were attached to LNPs containing either an active cGAMP or an inactive version of this (‘cGAMP Ctrl’).. (**B**) Tumor growth in mice administered either PBS or ADT alone, or these followed by administration every 2 days of the various forms of LNP listed. In panels **C** and **D**, various key groups have been selected from (**B**) and shown separately (for clarity). (**C**) Tumor growth curves in response to PBS alone (control) versus a single dose of ADT or (**D**) PBS alone (control), a single dose of ADT, ADT plus FRβ antibody-coated LNPs containing either cGAMP or cGAMP Ctrl. (**E**) PBS alone (no ADT) followed by FRβ antibody-coated LNPs containing either c-GAMP or cGAMP Ctrl. (**F**) ADT followed by Control IgG-coated LNPs containing either active cGAMP or cGAMP Ctrl. Data are presented as means ± SEMs. ***p□<□0.001 (comparing tumor sizes at sacrifice).

Mice bearing orthotopic Myc-CaP tumors were then administered either PBS alone (the vehicle for ADT) or a single dose of ADT alone on day 7 after implantation (‘ADT’ group). Two days later, separate groups of control or ADT-treated mice were administered the various forms of LNPs list in **Figure 3B** (see box). Controls included LNPs coated with no antibody or coated with a control rat IgG instead of the FRβ antibody, and others containing an inactive form of cGAMP (‘cGAMP Ctrl’) rather than active cGAMP. LNP injections were continued every two days until day 22, and tumor growth was assessed at regular intervals. None of the LNP groups appeared to have deleterious effects on the mice in terms of their eating/drinking behaviour, overall health and body weight (**Suppl. Fig. 6D**).

The effects of these various treatments on tumor growth are shown in **Figure 3B** (and selected groups are shown separately to facilitate comparisons in **Figure 3C-F**). A single dose of ADT was shown to have a similar effect on the growth of Myc-CaP tumors as two ADT doses, indicating the onset of CR within 7 days of ADT in this model (**Figures 1A & 3C**). The various control LNP groups showed no effect on tumor growth in the presence or absence of ADT (**Figure 3 E&F**). In contrast, LNPs coated with FRβ antibody and containing active cGAMP significantly delayed the onset of CR (**Figure 3D**).

When tumors from these mice were immunofluorescently stained, LNPs coated with FRβ antibody and containing active cGAMP (‘LNPs(E)’) were seen to be taken up mainly by PV F4/80+ TAMs rather than non-PV TAMs or F4/80- cells in PV or non-PV areas (**Figure 4A**). PV TAMs bearing LNPs were FRβ+ (**Figure 4B-D**) and FRβ antibody-coating of LNPs was essential for their uptake by PV TAMs. Only <5% of PV F4/80+ cells took up LNPs when they had either a control IgG or no IgG on their surface. This is supported by the fact that neither of these two LNP groups (with or without cGAMP) delayed the start of CR after ADT (**Figure 3b**).

**Figure 4.**
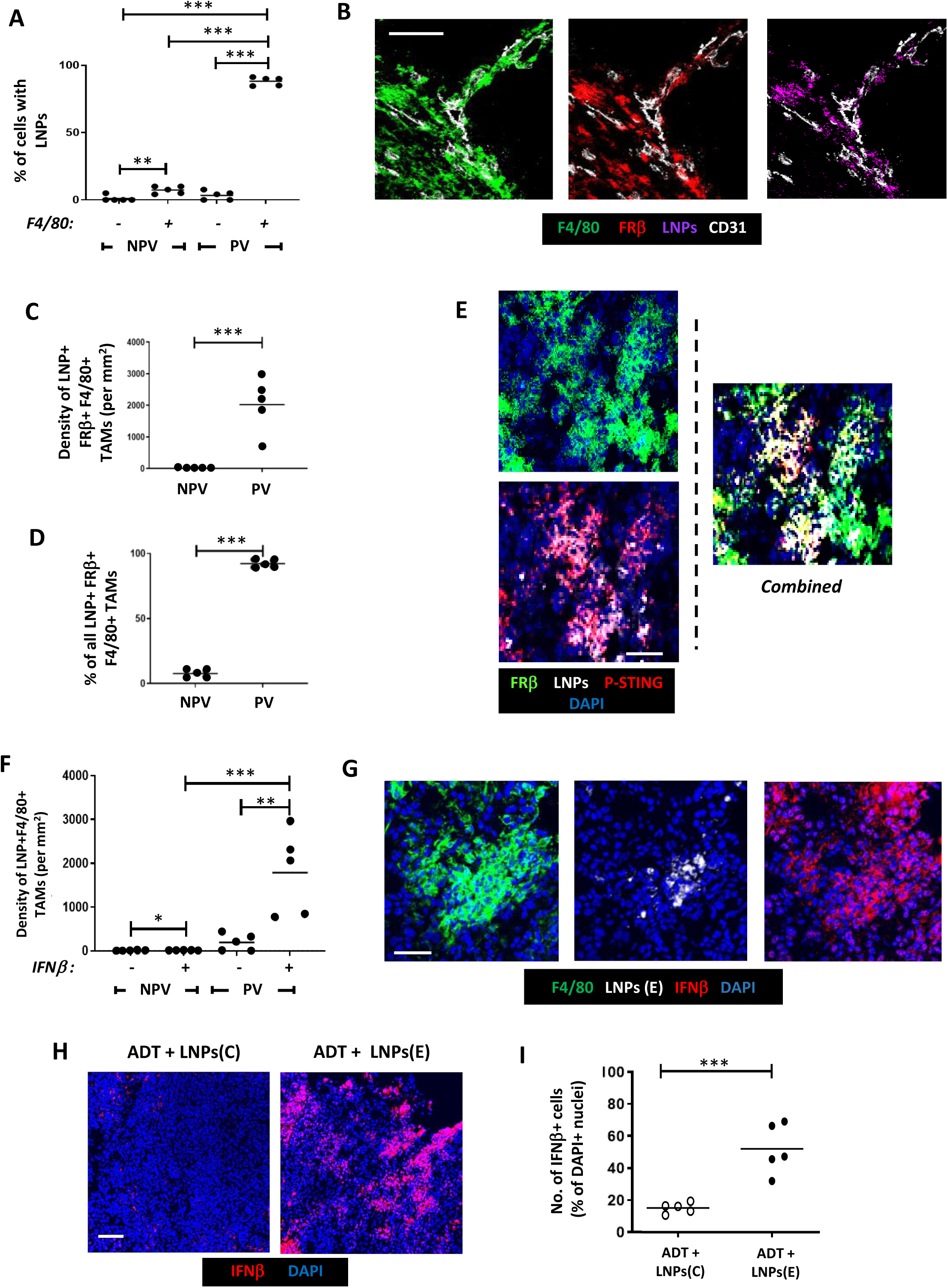
Selective delivery of cGAMP to PV FRβ+ TAMs in Myc-CaP tumors results in STING activation and upregulation of IFNβ. Following the administration of ADT plus LNPs: (**A**). The proportion of cells in PV and non-PV areas bearing LNPs that were F4/80- versus F4/80+. (**B**) Fluorescently labelled LNPs co-localised with PV FRβ+F4/80+ TAMs. (**C&D**) FRβ+F4/80+ TAMs bearing LNPs were only present in PV areas. (**E**) When LNPs bearing active cGAMP (LNPs(E)) were administered, the expression of active phosphorylated STING (P-STING) could be detected in LNP+FRβ+F4/80+ TAMs. This was accompanied by a significant increase in IFNβ detection PV LNP+F4/80+ TAMs (ie. in the LNPs(E) group) (**E**). In this group, IFNβ detection was only detectable in F4/80+ TAMs in PV not NPV areas, (**F**) but often extended beyond LNP+ cells, indicating its possible release and uptake by other cells in tumors (**G**). This did not occur when mice were injected with LNPs bearing inactive cGAMP. (NPV = non-PV). Data are presented as means ± SEMs. **p□<□0.01, and ***p□<□0.001. Magnification bars = 50µm.

Immunofluorescence analysis revealed that LNP+ FRβ+F4/80+ TAMs display p-STING in response to LNPs(E) treatment (**Figure 4E**). This was not seen with either FRβ+ antibody-coated LNPs containing inactive cGAMP (‘LNPs(C)’) or LNPs coated with control IgG (data not shown), indicating that the cGAMP-STING pathway was successfully activated only by LNPs(E) treatment.

The effect on cGAMP activation of STING on expression of IFNβ by PV FRβ+ TAMs was then demonstrated. The density of LNP+ TAMs expressing immunoreactive IFNβ was significantly higher in PV than non-PV areas of LNPs(E) - treated tumors. Indeed, very few IFNβ+LNP+ TAMs were present in non-PV areas of LNPs(E) -treated tumors (**Figure 4F**). Interestingly, IFNβ was detected beyond PV TAMs indicating the release of this cytokine by PV FRβ+ TAMs and subsequent uptake by neighbouring cells. This accords well with the finding that many cell types in tumors express receptors for type I IFNs.^33^ This increase in tumor IFNβ levels after LNPs(E) treatment was not observed with FRβ antibody-coated LNPs bearing the inactive form of cGAMP (‘LNPs(C)’) (**Figure 4H&I**), illustrating that LNP(E) treatment is specific and reliant on cGAMP-STING pathway activity.

We then examined the effect of elevated IFNβ (a known immunostimulant) on the density, distribution and activation status of CD8+ T cells, CD4+ T cells and NK cells. As shown in **Figure 2A** (after 2 doses of ADT), a single dose of ADT alone in the LNP experiment resulted in a significant increase in PV CD8+ T cells. Neither the co-administration of FRβ-antibody coated LNPs with LNPs(E) nor LNPs(C) with ADT altered this ADT-induced PV accumulation of CD8+ T cells (**Figure 5A&B**). However, ADT plus LNPs(E) increased the density and proportion of PV PD-1+CD8+ T cells (ie. reversed the induction by ADT alone of PV PD-1-CD8+T cells - see **Figure 2B**). This induction of active (PD-1+) CD8+ T cells by LNPs(E) occurred only in PV areas of ADT-treated tumors and did not occur with LNPs(C) (**Figure 5C**). Finally, we immunostained sections with antibodies against PD-1, LAG3 (a marker of T cell exhaustion) and CD8 to investigate the functional status of PV CD8+ T cells after LNP treatment. Fully active T cells were described as CD8+PD-1+LAG3- and exhausted ones as CD8+PD-1+LAG3+. **Figure 5F** shows that both PV and non-PV PD-1+CD8+ T cells in the ‘ADT plus LNPs(E)’ group were predominantly LAG3- (ie. fully active).

**Figure 5.**
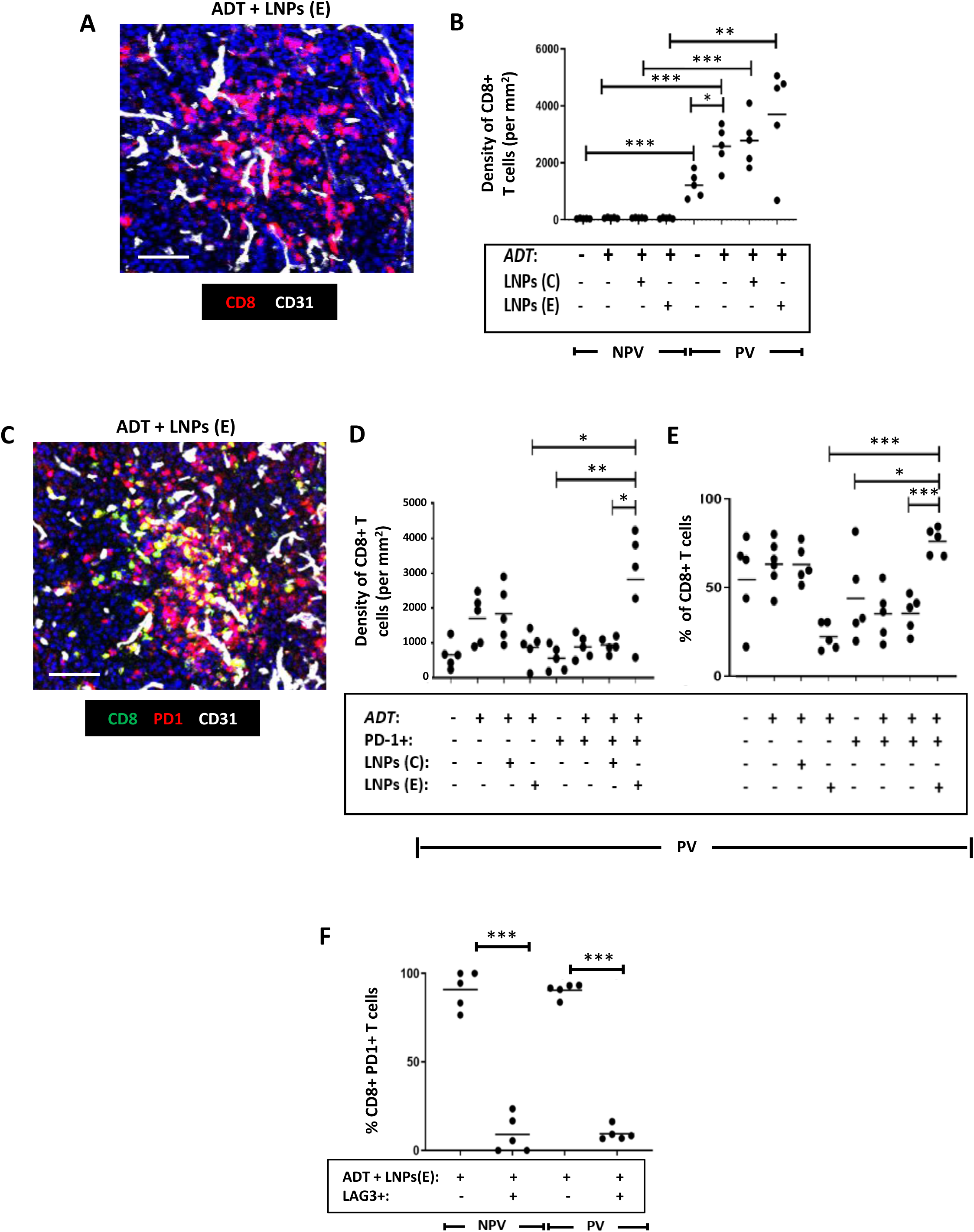
Selective delivery of cGAMP to PV FRβ+ TAMs in Myc-CaP tumors reverses the effect of ADT on the activation status of CD8+ T cells in Myc-CaP tumors. (**A**) Representative fluorescence image showing CD8+ T cells in a vascularised area of a tumor treated with ADT plus FRβ antibody-coated LNPs containing active cGAMP (‘LNPs(E)’). **(B)** ADT increased the PV accumulation of CD8+T cells (a change that was unaffected by co-administration with FRβ antibody-coated LNPs containing either inactive cGAMP (‘LNPs(C)’) or LNPs(E)). **(C)** Representative fluorescence image showing co-localization of PD-1 and CD8 in a vascularised area of a tumor treated with ADT plus LNPs(E). **(D,E)** ADT administered with LNPs(E) reversed the PV accumulation of inactive (PD-1-) CD8+ T cells induced by ADT alone, and led to an increase in both the PV density **(D)** and proportion **(E)** of CD8+ T cells expressing PD-1. **(F)** The majority of PD-1+CD8+ T cells did not express the exhaustion marker, LAG3 following treatment with ADT plus LNPs(E). (NPV = non-PV). Data are presented as means ± SEMs. *p□<□0.05, **p□<□0.01, and ***p□<□0.001. Magnification bars = 50µm.

Figure 6A&B show that CD4+ T cells are mainly PV in Myc-CaP tumors and remain so after ADT. There was a non-significant trend towards this PV accumulation of CD4+ T cells being increased further by the administration LNPs(E), but not LNPs(C) with ADT. Neither form of LNP altered the proportion of CD4+ T cells expressing the activation marker, PD-1, in PV areas but the density of PV PD-1+CD4+ T cells was significantly increased during ADT by LNPs(E), but not LNPs(C) (**Figure 6D&E**).

**Figure 6.**
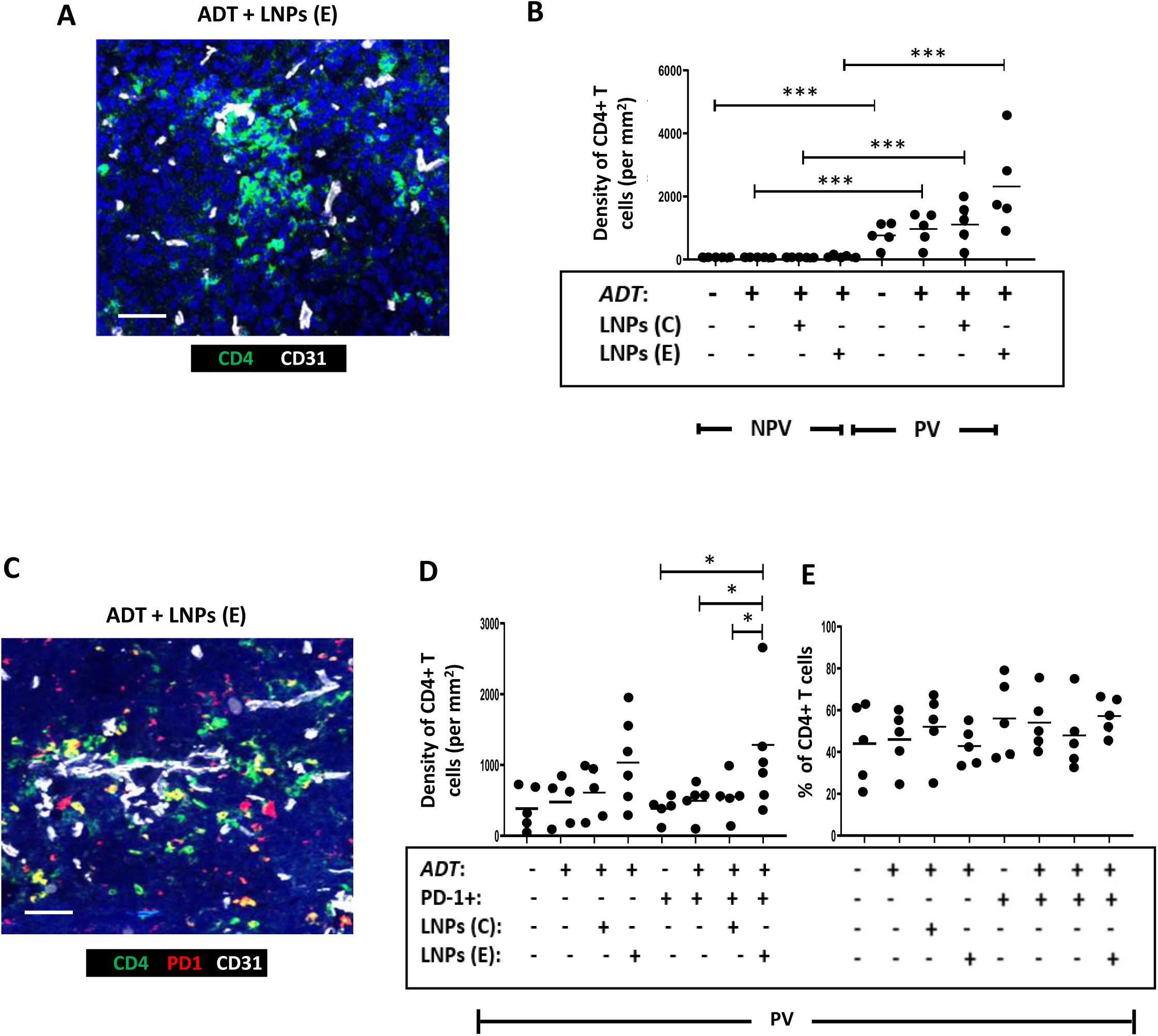
Selective delivery of cGAMP to PV FRβ+ TAMs in Myc-CaP tumors increased the PV accumulation of PD-1+CD4+ T cells in Myc-CaP tumors. (**A**) Representative fluorescence image showing CD4+ T cells in a vascularised area of a tumor treated with ADT plus FRβ antibody-coated LNPs containing active cGAMP (‘LNPs(E)’). (**B**) CD4+T cells were mainly PV in PBS-treated tumors and this was unaffected by ADT alone or co-administration of ADT with FRβ antibody-coated LNPs containing either inactive cGAMP (‘LNPs(C)’) or LNPs(E). **(C)** Representative fluorescence image showing co-localization of PD-1 and CD4 in a vascularised area of a tumor treated with ADT plus FRβ antibody-coated LNPs containing active cGAMP. **(D&E)** ADT administered with LNPs(E) resulted in the PV accumulation of PD-1+ CD4+ T cells. The proportion **of** CD4+ T cells expressing PD-1 remained unaltered by this treatment. (NPV = non-PV). Data are presented as means ± SEMs. *p□<□0.05 and ***p□<□0.001. Magnification bars = 50µm.

As mentioned previously, the density of NK cells is higher in PV than non-PV areas of both control and ADT-treated Myc-CaP tumors, and they mainly express CD69 in this location (**Suppl. Figure 5D**). This was not altered when LNPs(E) or LNPs(C) were administered with ADT (Figure 7), but the trend towards a lower density of activated PV NK cells after ADT alone (**Suppl. Figure 5D, left panel**) was reversed by LNPs(E) but not LNPs(C) (Figure 7D).

**Figure 7.**
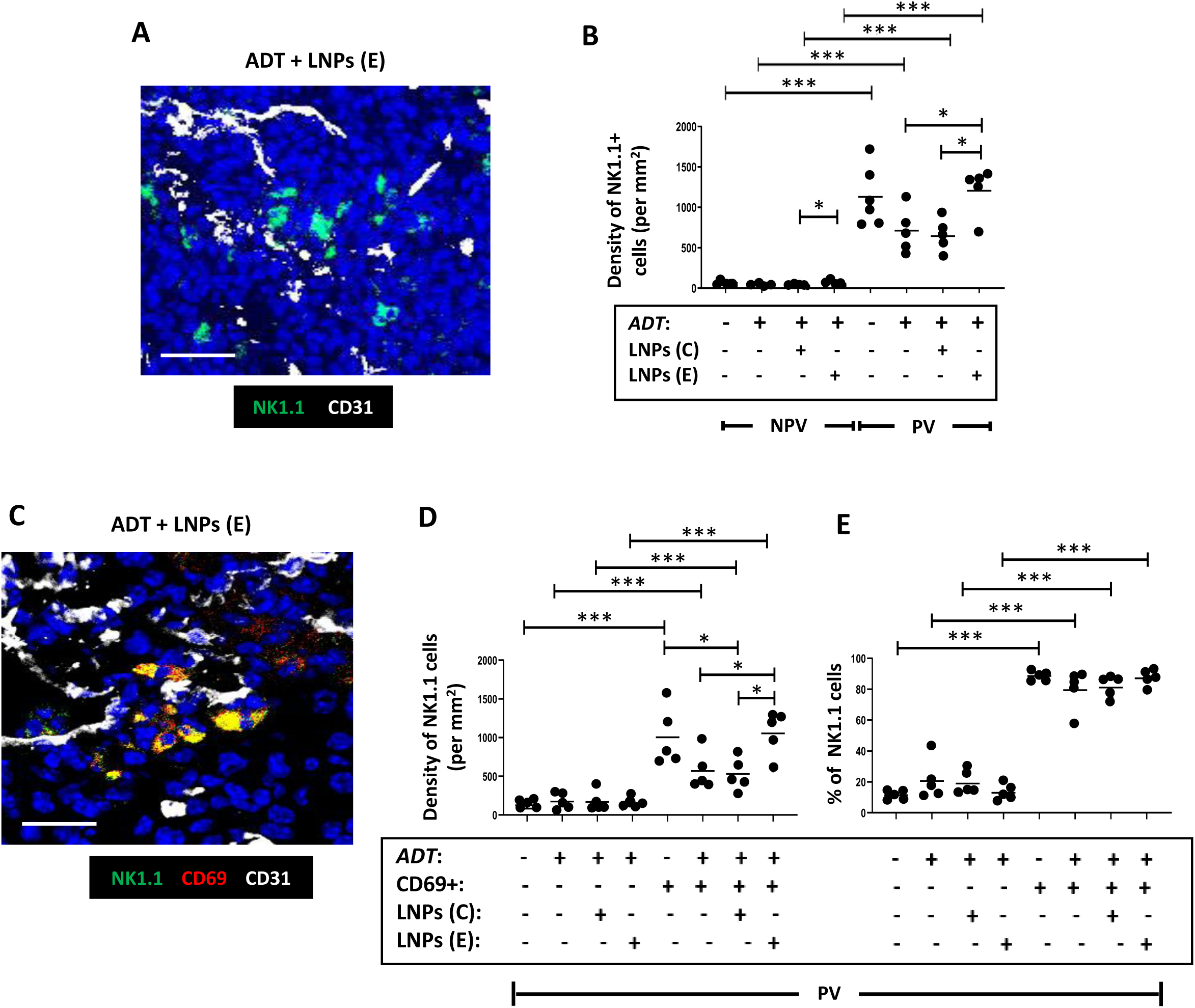
Selective delivery of cGAMP to PV FRβ+ TAMs in Myc-CaP tumors increases the PV density of active NK cells in Myc-CaP tumors. (**A**) Representative fluorescence image showing NK1.1+ NK cells in a vascularised area of a tumor treated with ADT plus FRβ antibody-coated LNPs containing active cGAMP (‘LNPs(E)’). (**B**) NK cells were mainly PV in PBS-treated tumors and this was unaffected by ADT alone. Co-administration of ADT with LNPs(E) increased both the non-PV and PV density of NK cells compared to ADT alone or ADT plus FRβ antibody-coated LNPs containing inactive cGAMP (‘LNPs(C)’). **(C)** Representative fluorescence image showing co-localization of CD69 and NK1.1 in a vascularised area of a tumor treated with ADT plus LNPs(E). **(D&E)** ADT administered with LNPs(E) increased the PV density of CD69+ (ie. activated) NK cells compared to ADT alone or ADT+ LNPs(C). The majority of PV NK cells expressed CD69 in PBS-treated tumors. This did not change after ADT, with or without LNPs. (NPV = non-PV). Data are presented as means ± SEMs. *p□<□0.05 and ***p□<□0.001. Magnification bars = 50µm.

The above findings indicate that when LNPs containing cGAMP are targeted to PV FRβ+ TAMs, this results in STING activation in these cells along with IFNβ release. This activated neighbouring immune cells like CD4+ and CD8+ T cells in the PV niche as well as the density of NK cells in these areas, and the resultant increase in anti-tumor immunity delays the onset of tumor regrowth (ie. the onset of CRPC) after ADT.

Finally, it was important to confirm that LNPs(E) do not target FRβ+ macrophages residing in healthy tissues. We, therefore, examined the effects of LNP exposure on a tissue known to contain FRβ+ macrophages - the liver (**Suppl. Figure 7**). FRβ was detected in >80% of F4/80+ Kupffer cells (**Suppl. Figure 7**). Despite this, only 20% of these cells took up LNPs or expressed IFNβ (**Suppl. Figure 7B&C**). When the extent of IFNβ immunoreactivity by all cells was examined in the liver, only 15-20% of all nucleated cells contained this cytokine (**Suppl. Figure 7D&E**). This matched the proportion of cells in the liver found to be FRβ+ F4/80+ Kupffer cells (**Suppl. Figure 7F**), and indicated that little, if any, IFNβ was taken up by other cell types in the liver.

## DISCUSSION

ADT is a mainstay treatment for prostate cancer but the development of CRPC limits its long-term efficacy in both the neoadjuvant and adjuvant settings.^1,32^ In the current study, we show that ADT induces the PV accumulation of both protumoral (FRβ+MRC1+CD169+VISTA+) TAMs in localised human and mouse prostate tumors prior to CRPC. Interestingly, TAMs with a similar phenotype accumulate in PV areas of mouse tumors during chemotherapy and promote tumor relapse.^33–35^ Further studies are now needed to confirm a causal link between such PV cells and tumor regrowth in CRPC.

ADT also causes naïve (PD-1-) CD8+T cells to gather in PV sites of prostate tumors suggesting their selective recruitment and/or suppression/retention in these areas. Protumoral PV TAMs are known to be immunosuppressive,^28,33,34^ so they may inhibit neighbouring T cells. A number of recent studies suggest this may be the case. For example, Bao and colleagues^36^ revealed a close cell-to-cell interaction between a subset of FRβ+MRC1+ TAMs and inactive (mainly PD-1-) CD8+ T cells in human colorectal carcinomas. Furthermore, depletion of FRβ+ TAMs expressing a transcriptional profile typical of PV TAMs (ie. *mrc1*, Lyve1, *Hmox1,Il10* and *stab1*) in a mouse model of ovarian cancer increased the frequency and activation of ascitic CD8+ T cells.^37^

Two proteins expressed by PV TAMs in ADT-treated tumors, VISTA and MRC1, could contribute to the inactive status of PV CD8+ T cells by binding to receptors/ligands on T cells to suppress their functions.^38,39^ This could inhibit T cell activation by dendritic cells as the latter are abundant in PV areas in mouse tumors^40^. CD169 was also expressed by PV TAMs which could also be immunosuppressive. A recent study revealed that CD169+ TAMs co-localize with T cells (along with regulatory T cells) in human breast tumors, and suppress T cells and NK cells, possibly via the release of PGE2, ROS and IL-1.^41^ Of particular relevance to the possible role of CD169+ TAMs in driving CRPC, is a recent study showing that CD169+ macrophages promote resistance to the anti-androgen, enzalutamide (an AR antagonist) in mouse bone metastases.^42^

The increase seen in PV naïve CD8+ T cells after ADT could also be due, in part, to thymus enlargement, a known side-effect of ADT.^43^ This enhances thymic release of naïve T cells into the circulation,^44^ which could lead to more then being recruited into tumors. However, thymic release of naïve CD4+ T cells also occurs during ADT,^45^ but these cells were not increased in ADT-treated mouse tumors. suggesting that other mechanisms are involved in the selective increase in naïve CD8+ T cells in PV areas.

The main aim of our study was to see whether ‘re-educating’ PV TAMs to enable them to release a potent immunostimulant, IFNβ, reverses the suppressive effects of ADT on tumor-infiltrating T cells. To do this, LNPs coated with an antibody to FRβ were used to deliver the STING agonist, cGAMP, to PV TAMs. In mouse prostate tumors, these LNPs were selectively taken up by PV TAMs after ADT, where they released cGAMP, activated STING signaling and the release IFNβ. The PV niche is an ideal location for this targeted form of immunotherapy as IFNβ has been shown to stimulate the functions of a number of immune cell types present in this site in tumors. For example, it stimulates antigen presentation by DCs to naïve T cells,^46^ as well as the proliferation and effector functions of T cells and NK cells.^47^ Type I IFNs can also decrease the immunosuppressive functions of MDSCs and Tregs and promote the expression of anti-tumor phenotype of TAMs.^48^ So, the anti-tumor effects of our LNP therapy would likely have been multifaceted in ADT-treated tumors and suggests that it could be used to enhance the efficacy of immunotherapy in prostate cancer. To date, clinical trials using immune checkpoint inhibitors, vaccines, CAR-T cells, and other forms of IT have shown limited efficacy in this disease.^49^

The FRβ-targeted delivery of cGAMP in LNPs to PV TAMs - and downstream IFNβ - could help circumvent the serious side effects recorded when non-encapsulated cGAMP or type I IFNs are injected into the systemic circulation. Indeed, no deleterious effects were seen in the livers of mice injected with cGAMP-containing LNPs despite the presence of FRβ-expressing Kupffer cells. The presence of LNPs and expression of IFNβ were detectable in only a small subset (20%) of these cells with no signs of IFNβ uptake by other cell types. This accords well with the finding that the expression of receptors for type I IFNs like IFNβ (IFNARs1 & 2) is negligible in the liver.^50^

Taken together, our studies show that LNP delivery of cGAMP to PV TAMs in ADT-treated prostate tumors increases anti-tumor immunity and delays CR. If this were to be reproduced in patients with intermediate/high-risk localised prostate tumors, it could extend the treatment window for neoadjuvant ADT and reduce the risk of progression to metastatic disease.

## DECLARATIONS

### Ethics approval and consent to participate

Studies with human tumor samples were approved by Research Ethics Committees at either the Institute of Cancer Research, Sutton, UK or the Dana Faber Cancer Institute in Boston, USA (as appropriate).

### Patient consent for publication of results

Written, informed consent was provided by all donors.

### Availability of data and material

Further information and requests for resources/reagents or data should be directed to and will be fulfilled by the lead contact, Claire Lewis.

### Competing interests

None declared.

### Funding

This project was funded by a grant from Prostate Cancer-UK to C.E.L. and J.E.B. (RIA16-ST2-022). HBP is supported by a CRUK Career Development award (#A27894).

### Contributors

C.E.L. and J.E.B. conceived the study and obtained funding for it. H.A-J., R.A., K.M., M.F., C.B. and H.P. performed experiments. H.A-J., K.M., M.C., B.G., H.N., C.B., A.J-J., M.M., H.P. and M.E-T. contributed to the experimental design and data analysis. C.E.L., J. DB., A.J-J., M.M., H.P., M.E-T. and J.E.B. supervised the study. C.E.L. drafted the original version of this manuscript, which was reviewed and edited by co-authors.

## Acknowledgements

We thank Dr Mukund Seshadri at the Roswell Park Comprehensive Cancer Center, USA, for the kind gift of the Myc-CaP cells stably transfected with luciferase.

## SUPPLEMENTARY FIGURE LEGENDS

**Suppl. Figure 1. PV F4/80+ TAMs also express CD169, VISTA and MRC1 in Myc-CaP tumors.** (**A**) Representative fluorescence images showing the co-localization of F4/80 with CD169, VISTA or MRC1 in vascularised tumor areas. Bar = 50µm. Quantitative analysis revealed the high density of TAMs expressing the following in PV areas after ADT: (**B**) F4/80 and CD169, (**C**) F4/80 and VISTA, and (**D**) F4/80 and MRC1. Data are presented as means ± SEMs. *p□<□0.05, **p□<□0.01, and ***p□<□0.001.

**Suppl. Figure 2. PV F4/80+ TAMs co-express CD169, VISTA and MRC1**

Quantitative analysis showed a similar proportion of F4/80+ TAMs expressing either: (**A**) CD169, (**B**) VISTA, and (**C**) MRC1 in both PV and non-PV areas of tumors. Data are presented as means ± SEMs. *p□<□0.05, **p□<□0.01, and ***p□<□0.001. (**D**) Representative fluorescence images showing the co-localization of F4/80 with CD169, VISTA and MRC1 in a vascularised tumor area. Bar = 50µm.

**Suppl. Figure 3. Effect of physical castration on TAM distribution in transgenic mouse prostate (Ptenfl/fl) tumors.** The density (**A**) or proportion (**B**) F4/80+ TAMs expressing MRC1 in PV and non-PV (NPV) areas of tumors in sham-castrated or castrated mice. Data are presented as means ± SEMs. *p□<□0.05 and **p□<□0.01. (**C**) The majority of PV F4/80+ TAMs in ADT-treated tumors also expressed FRβ. Bar = 50µm.

**Suppl. Figure 4. FRβ expression by PV CD68+TAMs in localised human prostate tumors before and after ADT: correlation with tumor response.** (**A**) CD68+ TAMs and (**B**) FRβ+CD68+ TAMs were predominantly PV both before and after patients received ADT (ie. matching samples). Tumors showing signs of CR (ie. starting to regrow within six months of ADT treatment - termed ‘non-responders’) were also compared to those that were still responsive to ADT (ie. hormone sensitive, ‘HS’ – termed ‘responders’), Data are presented as means ± SEMs. *p□<□0.05 and ***p□<□0.001. ‘ns’ = not significant.

**Suppl. Figure 5. Effect of ADT on the distribution and activation status of CD4>+ T cells and NK cells in Myc-CaP tumors.** (**A**) Representative fluorescence image of the co-localization of CD4 and PD-1 in a vascularised area of an ADT-treated tumor. (**B**) Density (left panel) and proportion of CD4+ T cells expressing PD-1 in PV and non-PV (NPV) areas. (**C**) Representative fluorescence image of the co-localization of NK1.1 and CD69 in a vascularised area of an ADT-treated tumor. (**D**) Density (left panel) and proportion (right panel) of NK cells expressing CD69 in PV and non-PV (NPV) areas. Data are presented as means ± SEMs. *p□<□0.05, **p□<□0.01, and ***p□<□0.001. Bars = 50µm.

**Suppl. Figure 6. Rationale and design of antibody-coated LNPs to target PV FRb+ TAMs in ADT-treated Myc-CaP tumors.** (**A**) Schematic illustration of the methodology used to generate antibody-coated LNPs containing either inactive or active cGAMP. Abbreviations used: MC3, D-Lin-MC3-DMA. DSPC, 1,2-distearoyl-sn-glycero-3-phosphocholine. Chol, cholesterol.DSPE-PEG2000, 1,2-distearoyl-sn-glycero-3-phosphoethanolamine-N-[methoxy(polyethylene glycol)-2000]. OG488-DHPE, Oregon Green 488 conjugated 1,2-dihexadecanoyl-sn-glycero-3-phosphoethanolamine.DSPE-PEG2000-DBCO,1,2-distearoyl-sn-glycero-3-phosphoethanolamine-N-[dibenzocyclooctyl (polyethylene glycol)-2000] DBCO, dibenzocyclooctyne. cGAMP, 2’3’-cyclic guanosine monophosphate–adenosine monophosphate. LNP, lipid nanoparticle. N3, azide. FRβ, folate receptor beta. UDP-GalNAz, UDP-N-azidoacetylgalactosamine. (**B**) Flow cytometry showing the expression of FRα but not FRβ by Myc-CaP cells *in vitro*. (**C**) Representative fluorescence images of (**left panel**) FRα+ cancer cells (green) and FRβ staining (red) on separate cell populations. TCI = tumor cell islands. FRβ staining of F4/80+ TAMs (yellow in **right panel**). ***p□<□0.001. Magnification bar = 50µm. **(D)** Neither ADT no ADT plus LNPs coated with FRβ antibody (containing either active or inactive cGAMP) altered mouse body weight.

**Suppl. Fig. 7. Minimal expression of IFNβ in the liver of mice bearing Myc-CaP tumors following administration of ADT plus LNPs.** (**A**) The majority (>80%) of F4/80+ Kupffer cells in the liver expressed FRβ in mice administered ADT plus LNPs containing either active (LNPs(E)) or inactive (LNPs(E)) cGAMP. However, LNPs (**B**) and IFNβ (**white arrows in C)** were only detected in approx. 20% of FRβ+ cells. Overall, IFNβ staining was detected in 10-20% of all nucleated cells in the liver of mice administered ADT plus LNPs(E) (**D&E**). This matched the proportion of all nucleated cells in the liver that were FRβ+F4/80+ macrophages (**F**). Data are presented as means ± SEMs. *p□<□0.05, **p□<□0.01, and ***p□<□0.001. Bars = 50µm.

**Suppl. Table. Clinicopathological data for the anonymised patient groups whose tumor sections were stained and analysed in Figures 2 and 3.** These were supplied by the Institute of Cancer Research (ICR) in London, UK and the Dana Faber Cancer Institute in Boston, USA. Abbreviations used: ADT, androgen deprivation therapy; RP, radical prostatectomy; TURP, Transurethral resection of the prostate, LHRH, luteinising hormone-releasing hormone.

## Notes

### Competing Interest Statement

The authors have declared no competing interest.

