## Supplementary figures and images for "Targeting of a STING Agonist to Perivascular Macrophages in Prostate Tumors Delays Resistance to Androgen Deprivation Therapy"

### Suppl. Figures

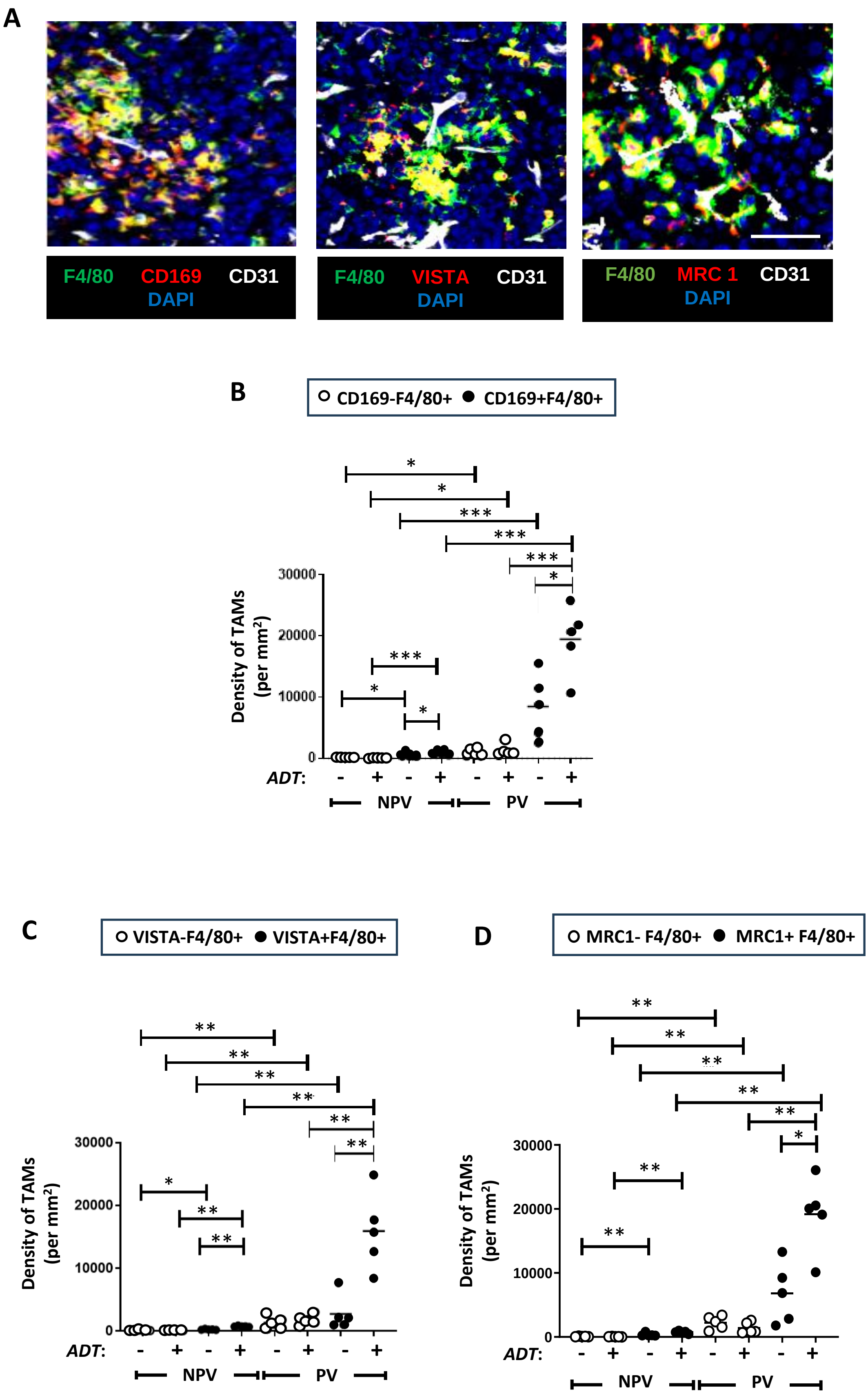

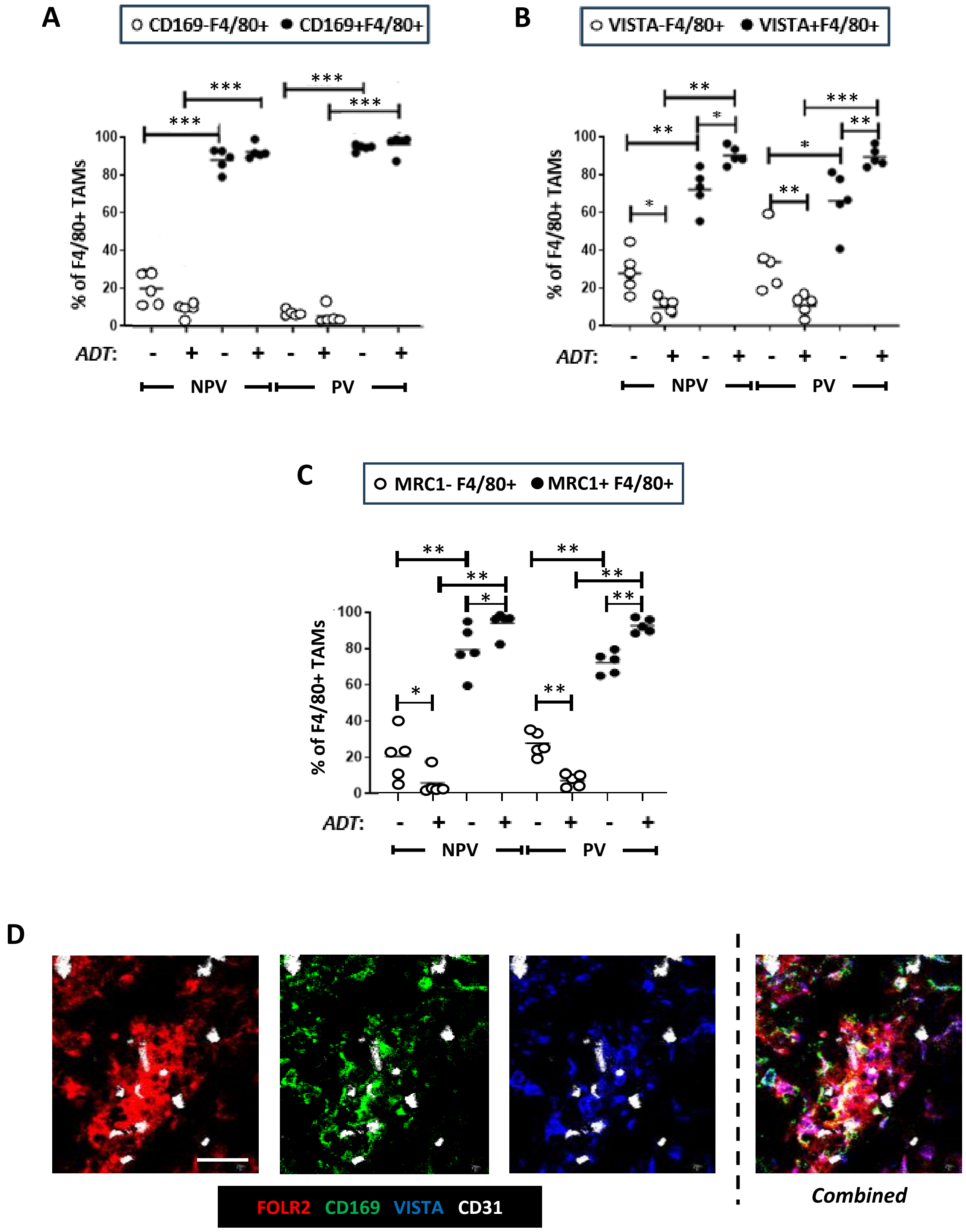

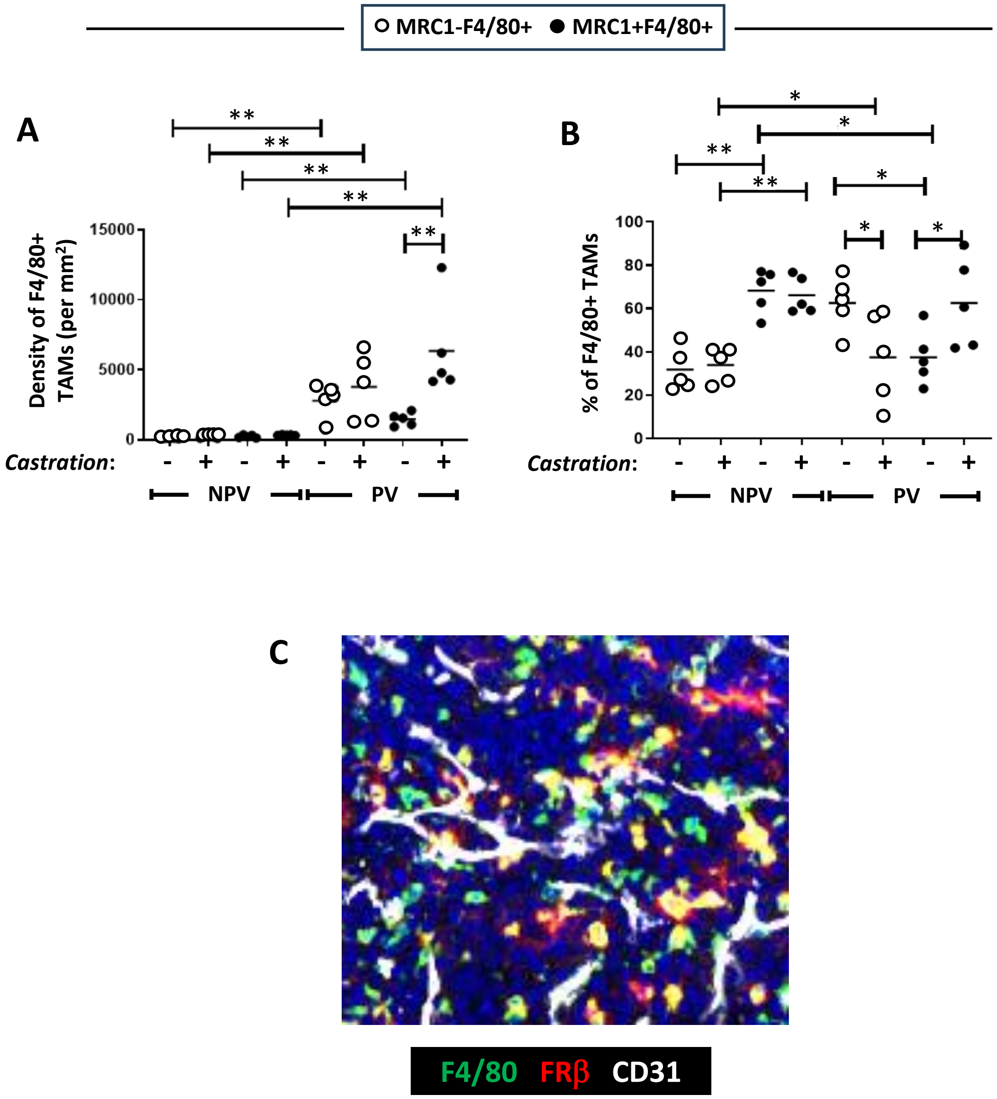

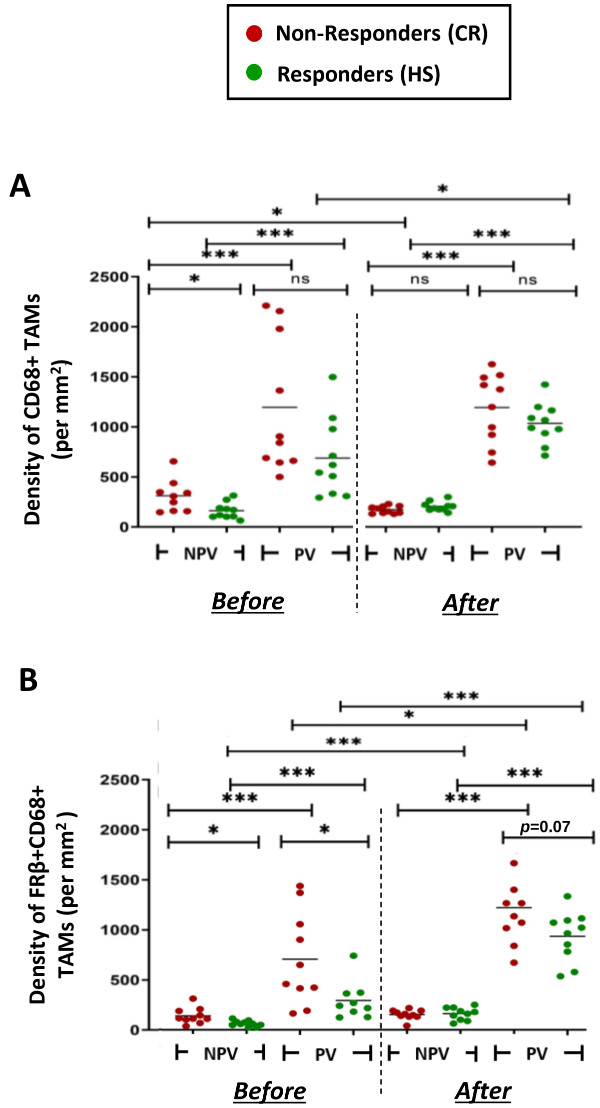

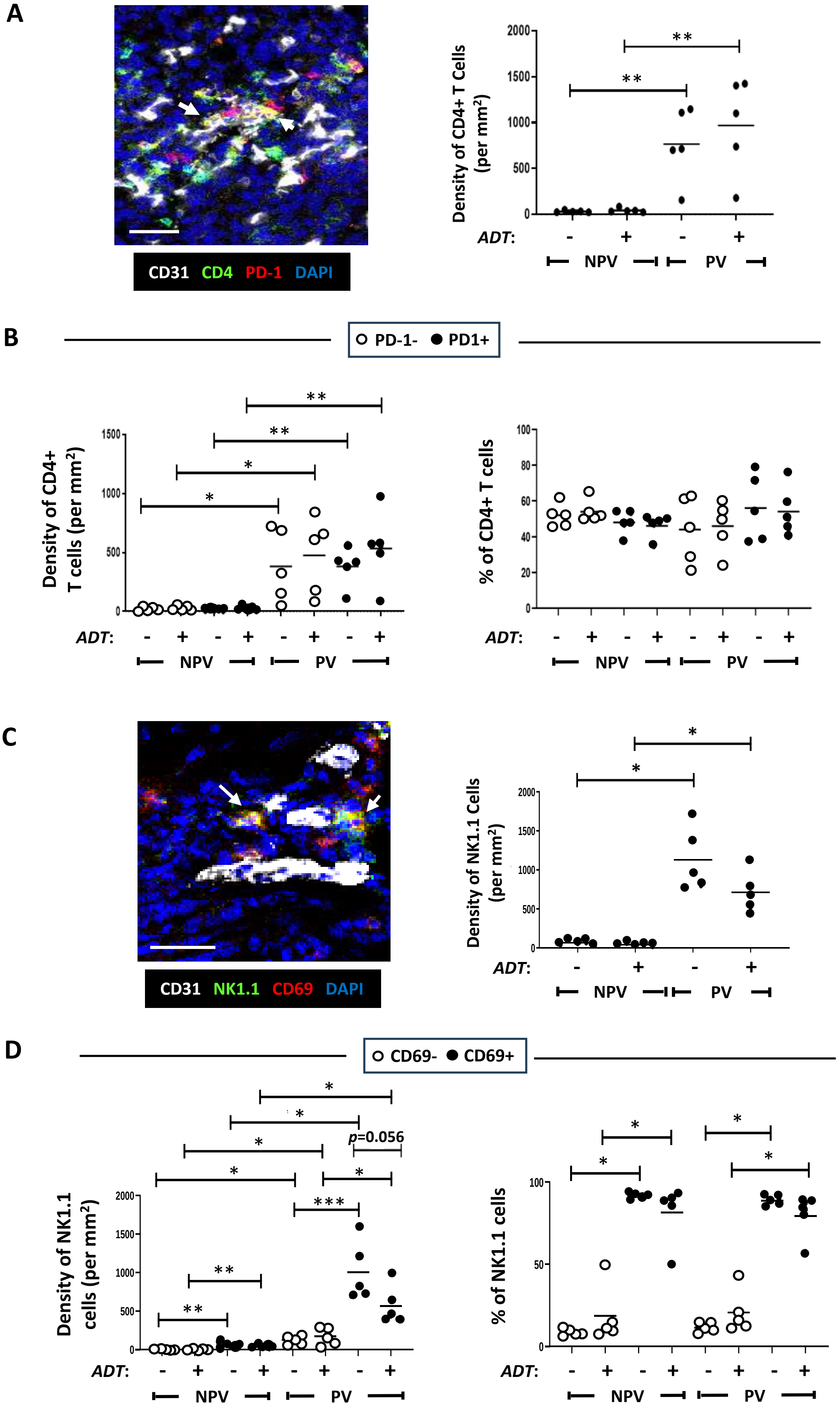


Suppl. Figure 5

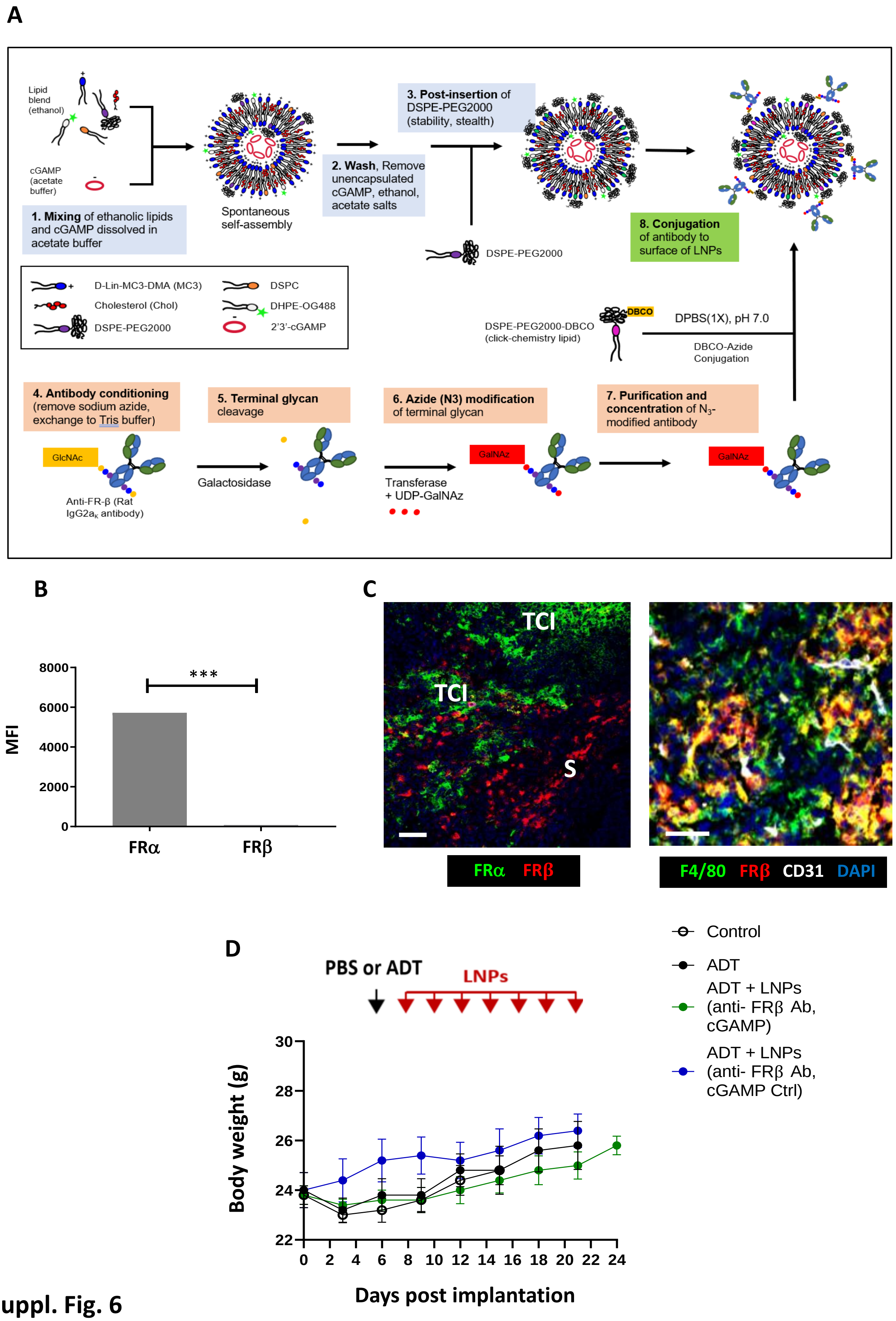

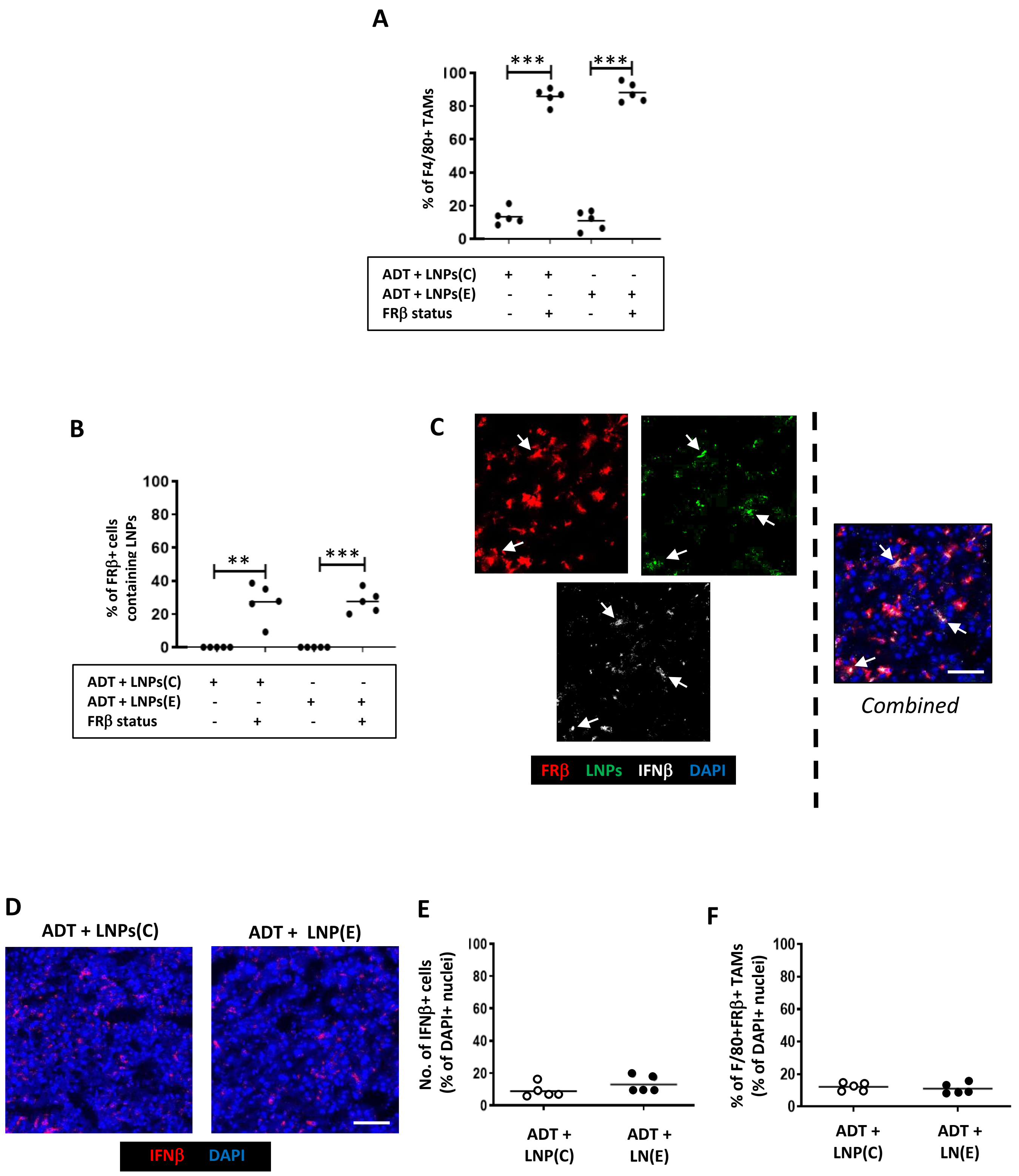
