## Supplementary material for "Targeting of a STING Agonist to Perivascular Macrophages in Prostate Tumors Delays Resistance to Androgen Deprivation Therapy": Suppl. Table

| | London - (untreated; No ADT) | London – ADT treated | Harvard Non-responders | Harvard Responders |
|---|---|---|---|---|
| **Age at diagnosis** | | | | |
| 40-59 | 2 | 2 | 5 | 4 |
| 60-79 | 3 | 3 | 5 | 5 |
| 80-99 | 1 | 0 | 0 | 0 |
| **Gleason score at RP/TURP** | | | | |
| 6 | 1 | 0 | 0 | 0 |
| 7 | 2 | 1 | 1 | 2 |
| 8 | 0 | 1 | 4 | 2 |
| 9 | 3 | 2 | 5 | 5 |
| 10 | 0 | 1 | 0 | 1 |
| **Stage at RP/TURP** | | | | |
| T0 | 0 | 0 | 0 | 5 |
| T1 | 0 | 0 | 0 | 0 |
| T2 | 1 | 0 | 0 | 5 |
| T3a | 2 | 0 | 6 | 0 |
| T3b | 1 | 2 | 4 | 0 |
| T4 | 0 | 2 | 0 | 0 |
| UNK | 2 | 1 | | |
| **Metastatic at RP/TURP:** | | | | |
| Yes | 0 | 3 | 0 | 0 |
| No | 6 | 2 | 0 | 0 |
| <u>**Treatments**</u> | | | | |
| ***ICR*:** | | | | |
| LHRH inhibitors (Leuprolide, Goserelin or Triptorelin) | 6 | 5 | - | - |
| ***Dana Faber Cancer Institute*:** | | | | |
| Group 1: Abiraterone + Predisone + Enzalutamide + Leuprolide | - | - | 3 | 3 |
| Group 2: Enzalutamide + Leuprolide | - | - | 3 | 1 |
| Group 3: Abiraterone + Predisone + Apalutamide + Leuprodide | - | | 1 | 2 |
| Group: Abiraterone + Predisone + Leuprolide | - | - | 3 | 4 |
